# A subclass of evening cells promotes the switch from arousal to sleep at dusk

**DOI:** 10.1101/2023.08.28.555147

**Authors:** Matthew P. Brown, Shubha Verma, Isabelle Palmer, Adler Guerrero Zuniga, Clark Rosensweig, Mehmet F. Keles, Mark N. Wu

## Abstract

Animals exhibit rhythmic patterns of behavior that are shaped by an internal circadian clock and the external environment. While light intensity varies across the day, there are particularly robust differences at twilight (dawn/dusk). These periods are also associated with major changes in behavioral states, such as the transition from arousal to sleep. However, the neural mechanisms by which time and environmental conditions promote these behavioral transitions are poorly defined. Here, we show that the E1 subclass of *Drosophila* evening clock neurons promotes the transition from arousal to sleep at dusk. We first demonstrate that the cell-autonomous clocks of E2 neurons alone are required to drive and adjust the phase of evening anticipation, the canonical behavior associated with “evening” clock neurons. We next show that conditionally silencing E1 neurons causes a significant delay in sleep onset after dusk. However, rather than simply promoting sleep, activating E1 neurons produces time- and light- dependent effects on behavior. Activation of E1 neurons has no effect early in the day, but then triggers arousal before dusk and induces sleep after dusk. Strikingly, these phenotypes critically depend on the presence of light during the day. Despite their influence on behavior around dusk, *in vivo* voltage imaging of E1 neurons reveals that their spiking rate does not vary between dawn and dusk. Moreover, E1-specific clock ablation has no effect on arousal or sleep. Thus, we suggest that, rather than specifying “evening” time, E1 neurons act, in concert with other rhythmic neurons, to promote behavioral transitions at dusk.

## INTRODUCTION

Daily rhythms of behavior depend on both timing information from internal circadian clocks and predictable changes in the external environment. In particular, the rising and setting of the sun produce dramatic changes in environmental light at dawn and dusk, respectively. Given the ethological importance of these transitions between daytime and nighttime, discrete changes in behavioral state often occur around these times, such as the switch from wakefulness to sleep. However, while much is known about how environmental cues, such as light, entrain the circadian clock, it remains unclear whether and how the clock interacts with these environmental changes at twilight to produce specific patterns of behavior.

Circadian rhythms are centrally controlled by groups of pacemaker neurons in the brain that rhythmically express core clock genes.^1^ In *Drosophila*, the central clock network is comprised of approximately 75 pacemaker neurons in each brain hemisphere.^2^ The clock network promotes robust rhythms of locomotor activity and sleep that demonstrate the ability of a fly to both predict and react to changing environmental conditions. As crepuscular animals, flies exhibit two peaks of activity, one occurring at dawn and one occurring at dusk. These two peaks of activity are thought to be separately controlled by different groups of lateral neurons (LNs) in the clock network: “morning cells” (the PDF-positive small ventral LNs/s-LNvs) and “evening cells” (E cells; the PDF-negative 5^th^ s-LNv and dorsal LNs/LNds).^3–7^ *Drosophila* exhibit an increase in sleep during midday (“siesta”), but, as dusk approaches, flies slowly increase their activity in anticipation of the light-to-dark transition. After dusk, they react to nightfall by rapidly transitioning from peak activity into sleep.^8–10^ A subset of dorsal clock neurons (DN1ps) has been shown to contribute to evening anticipatory behavior at the end of the “siesta”.^7^ However, most work on evening anticipation has focused on the E cells, as it has previously been shown that the clocks within these neurons are largely responsible for driving evening anticipation.^3,4,7^ Additionally, the clocks of the E cells can dictate the phase of activity in the evening,^5,6,11^ and the LNds exhibit peak levels of intracellular calcium slightly before dusk, further supporting their role in controlling evening behavior.^11–13^

The E cells are often referred to as a single, homogenous oscillator. However, genetic, morphological, and physiological evidence suggest the E cells can be further divided into three distinct subclasses, called E1, E2, and E3.^14–18^ Despite the differences among the subclasses of evening lateral neurons, most prior studies of these neurons have manipulated both E1 and E2 neurons together, potentially obscuring the individual contributions of these subclasses to behavior.^3,4,6,11,14,19,20^ We hypothesized that the E1 and E2 subclasses regulate distinct aspects of evening behavior around dusk.

In this study, we used subclass-specific genetic tools to selectively manipulate E1 and E2 subclasses of E cells and then assessed changes to daily, rhythmic locomotor behavior and sleep. We found that the cell-autonomous clocks of E2 neurons are primarily responsible for the previously described role of the E LNs in evening anticipation, while the clocks of E1 neurons are dispensable for this behavior. Instead, our data suggest that E1 neurons control the transition from arousal to sleep at dusk. Conditional silencing of E1 neurons causes a significant delay in sleep onset time, while conditionally activating E1 neurons has time- and light-dependent effects on arousal that are centered around dusk. Remarkably, E1 neuron activation has no effect early in the day, but can promote both activity and sleep around dusk in a light-dependent manner. Interestingly, we also found that the electrical activity of E1 neurons does not cycle between dawn and dusk *in vivo* and that the timing of their behavioral output does not depend on their own, cell-autonomous clocks. These results demonstrate that E2 neurons exhibit the canonical features of evening cells, while the E1 neurons are atypical clock neurons that play a newly defined role in promoting the behavioral transition from wakefulness to sleep at dusk.

## RESULTS

### E2 neuron clocks drive evening anticipation

Most prior work investigating the function of evening cells has focused on manipulating the core molecular clock within these cells while monitoring changes in rhythmic locomotor behavior. However, these studies have mostly used genetic driver lines that label multiple subclasses of evening cells.^3–6,11,19–21^ We hypothesized that evening anticipatory behavior is attributable to the E2 neurons, because the broader drivers previously used typically included these cells (E1 + E2 neurons in *Mai179-GAL4*^22^ and *MB122B-split GAL4*^20^ or E2 + E3 neurons in *dvpdf-GAL4*,^23^ **Fig. 1A**). To test whether E2 neurons are responsible for the previously observed effects on anticipation, we compared core clock manipulations in a combined E1+E2 driver, *R78G02-GAL4*^24^ (**Fig. 1B**) against an E2-specific driver, *R54D11-GAL4*^17,25^ (**Fig. 1C**). Using *R78G02-GAL4*, we ablated core clock rhythms in both E1 and E2 neurons through tissue-specific knockout of the core clock gene *timeless* (*tim*) using a previously-validated guide RNAs and assessed daily locomotor behavior in 12:12 light:dark (LD) cycles.^26^ As expected, this manipulation significantly reduced evening anticipatory behavior, highlighting the necessity of the molecular clock in evening cells for normal evening anticipatory behavior (**Figs. 1D and 1E**). In addition, changing the period length of molecular clocks in the evening cells has been shown to alter the phase of activity in the evening.^5–7,11,27^ Indeed, we also found that shortening the period length of the core clocks in both E1 and E2 neurons using the hyperactive kinase DBT^S^ advanced the phase of evening activity (**Figs. 1F and 1G**),^28^ further emphasizing the importance of the molecular clock in the evening cells for proper evening behavior.

**Figure 1.**
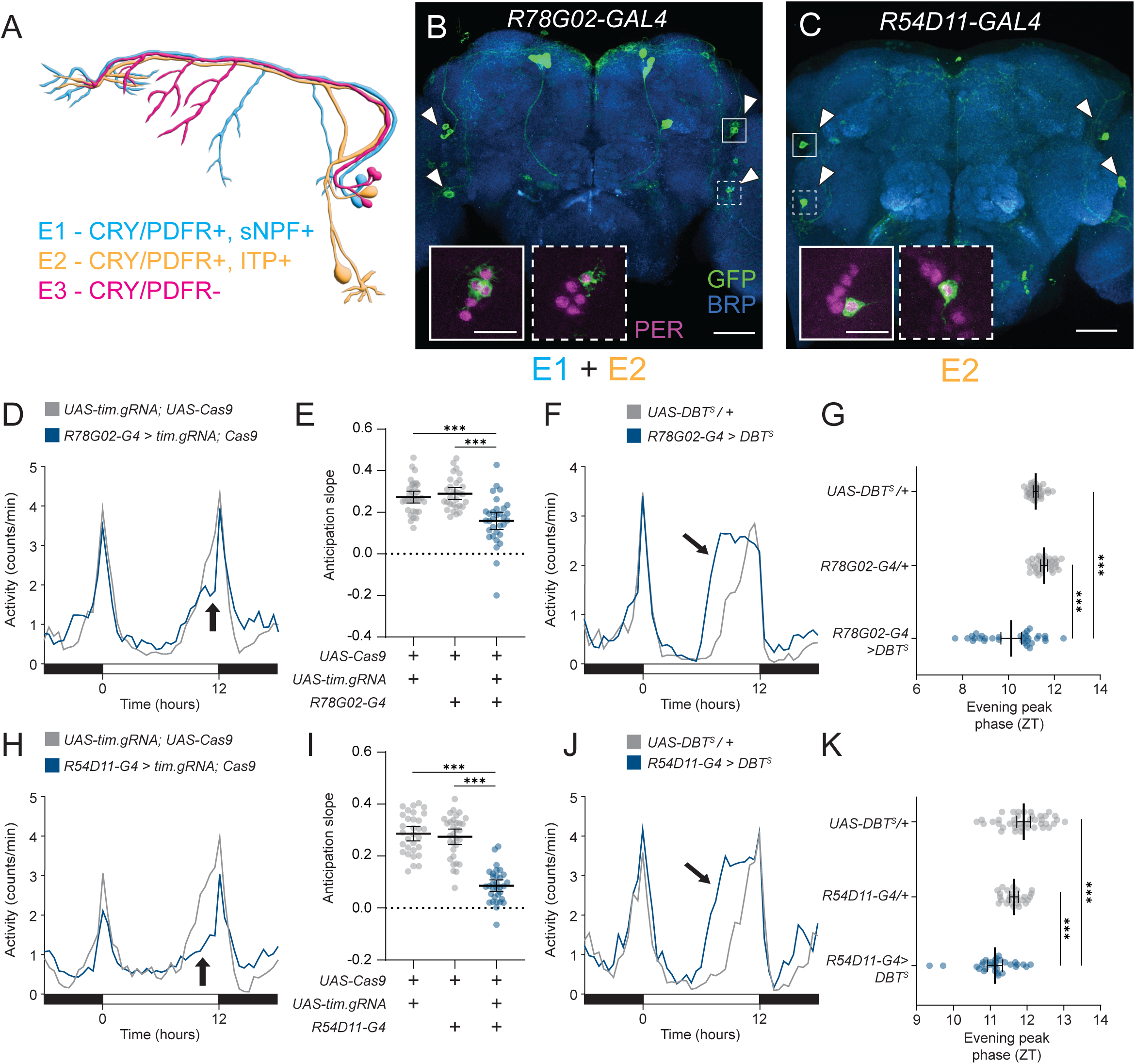
The core clock in E2 neurons drives evening anticipation. **(A)** Schematic diagram of evening cell subtypes and characteristic gene expression. CRY = Cryptochrome, PDFR = PDF Receptor, sNPF = small Neuropeptide F, ITP = Ion Transport Peptide. **(B and C)** Immunostaining of E cell drivers. Whole-mount immunostaining of *R78G02-GAL4* **(B)** and *R54D11-GAL4* **(C)** brains expressing *UAS-CD8::GFP* with anti-GFP (green), anti-Bruchpilot (BRP, blue), and anti-Period (PER, magenta, inset) antibodies. Target E cells indicated by arrowheads. PER staining highlighted in solid white inset (LNds) and dotted line inset (5^th^ s-LNv). Scale bars, 50 µm (main) and 15 µm (inset). **(D)** Tissue-specific knockout of *timeless* in both E1 and E2 neurons. Activity counts per min profile for 24 hrs of *R78G02-GAL4>UAS-tim.gRNA, UAS-Cas9.P2* (blue) and *UAS-tim.gRNA/+; UAS-Cas9.P2/+* (gray) flies in 12:12 LD plotted in 30 min bins. Arrow indicates altered evening anticipation. **(E)** Average slope of evening anticipation (see Methods) for *R78G02-GAL4>UAS-tim.gRNA, UAS-Cas9.P2* (blue, n=32) flies compared to *UAS-tim.gRNA/+; UAS-Cas9.P2/+* (gray, n=32) and *R78G02-GAL4>UAS-Cas9.P2* (gray, n=30) flies. Mean ± 95% confidence interval is shown. One-way ANOVA with Bonferroni post hoc test. **(F)** DBT^S^ expression in both E1 and E2 neurons. Activity counts per min profile for 24 hrs of *R78G02-GAL4>UAS-DBT^S^* (blue) and *UAS-DBT^S^/+* (gray) flies in 12:12 LD plotted in 30 min bins. Arrow indicates advanced evening peak. **(G)** Average phase of peak evening activity (right) for *R78G02-GAL4>UAS-DBT^S^* (blue, n=32) flies compared to *UAS-DBT^S^/+* (gray, n=32) and *R78G02-GAL4/+* (gray, n=32) flies. Mean ± 95% confidence interval is shown. One-way ANOVA with Bonferroni post hoc test. **(H)** Tissue-specific knockout of *timeless* specifically in E2 neurons. Activity counts per min profile for 24 hrs of *R54D11-GAL4>UAS-tim.gRNA, UAS-Cas9.P2* (blue) and *UAS-tim.gRNA/+; UAS-Cas9.P2/+* (gray) 12:12 LD plotted in 30 min bins. Arrow indicates altered evening anticipation. **(I)** Average slope of evening anticipation (see Methods) for *R54D11-GAL4>UAS-tim.gRNA, UAS-Cas9.P2* (blue, n=32) flies compared to *UAS-tim.gRNA/+; UAS-Cas9.P2/+* (gray, n=31) and *R54D11-GAL4>UAS-Cas9.P2* (gray, n=32) flies. Mean ± 95% confidence interval is shown. One-way ANOVA with Bonferroni post hoc test. **(J)** DBT^S^ overexpression specifically in E2 neurons. Activity counts per min profile for 24 hrs of *R54D11-GAL4>UAS-DBT^S^* (blue) and *UAS-DBT^S^/+* (gray) flies in 12:12 LD plotted in 30 min bins. Arrow indicates advanced evening peak. **(K)** Average phase of peak evening activity (right) for *R54D11-GAL4>UAS-DBT^S^* (blue, n=31) flies compared to *UAS-DBT^S^/+* (gray, n=40) and *R54D11-GAL4/+* (gray, n=26) flies. Mean ± 95% confidence interval is shown. One-way ANOVA with Bonferroni post hoc test. In this and in subsequent figures: *, *P* < 0.05; **, *P* < 0.01; ***, *P* < 0.001; ns, not significant. White and black bars indicate light and dark periods, respectively.

Interestingly, knocking out *tim* in E2 neurons alone using *R54D11-GAL4* similarly reduced evening anticipation (**Figs. 1H and 1I**), phenocopying the effect observed with *tim* knockout in both E1 and E2 neurons. In addition, overexpression of DBT^S^ using *R54D11-GAL4* significantly advanced the phase of evening activity (**Figs. 1J and 1K**), again reproducing the phenotype seen with DBT^S^ overexpression in both E1 and E2 neurons. Thus, manipulating the molecular clocks of E2 neurons alone is sufficient to produce the effects on evening activity previously described when manipulating multiple subclasses of evening cells.

One model for clock neuron function postulates that the cell-autonomous clock within each clock neuron drives rhythmic electrical activity to drive cyclical behavior. To test whether E2 neurons regulate evening behavior through their electrical activity, we conditionally silenced E2 neurons using the inwardly rectifying K^+^ channel Kir2.1 in combination with the auxin-inducible GAL4 expression system (AGES).^29,30^ Unexpectedly, auxin-induced silencing of E2 neurons with Kir2.1 did not impair evening anticipation, compared to controls (**Fig. S1A and S1B**). Next, we activated E2 neurons using the heat-sensitive cation channel dTrpA1.^31^ Similarly, thermogenetic activation of E2 neurons had no significant effect on evening anticipation, although it did lead to a significant increase in nighttime activity (**Figs. S1C-S1E**). Thus, these data suggest that E2 neuron control of evening anticipation is clock-dependent, but not activity-dependent.

### E1 neural activity promotes arousal before dusk and sleep onset after dusk

Since the E2 neurons have a strong influence on evening anticipatory behavior, we next asked what role E1 neurons play in controlling rhythmic, daily activity. To address this question, we used two GAL4 drivers that selectively express in E1 neurons, *Trissin-GAL4*^15,32^ and *R15C11-GAL4.*^33^ These GAL4 drivers have non-overlapping off-target expression, and among clock network neurons, *Trissin-GAL4* and *R15C11-GAL4* only express in E1 LNds (**Fig. 2A**). Interestingly, ablating clock rhythms in E1 neurons using *tim* CRISPR knockout had no effect on evening anticipation, suggesting that, unlike E2 molecular clocks, E1 molecular clocks are not necessary for evening anticipation (**Figs. S2A and S2B**). In addition, shortening the period length in E1 neurons using *UAS-DBT^S^*had only a modest effect with *R15C11-GAL4* and inconsistent effects with *Trissin-GAL4* (**Figs. S2C and S2D**), suggesting that the clocks in E1 neurons play a less important role than E2 neurons in setting the phase of evening activity.

**Figure 2.**
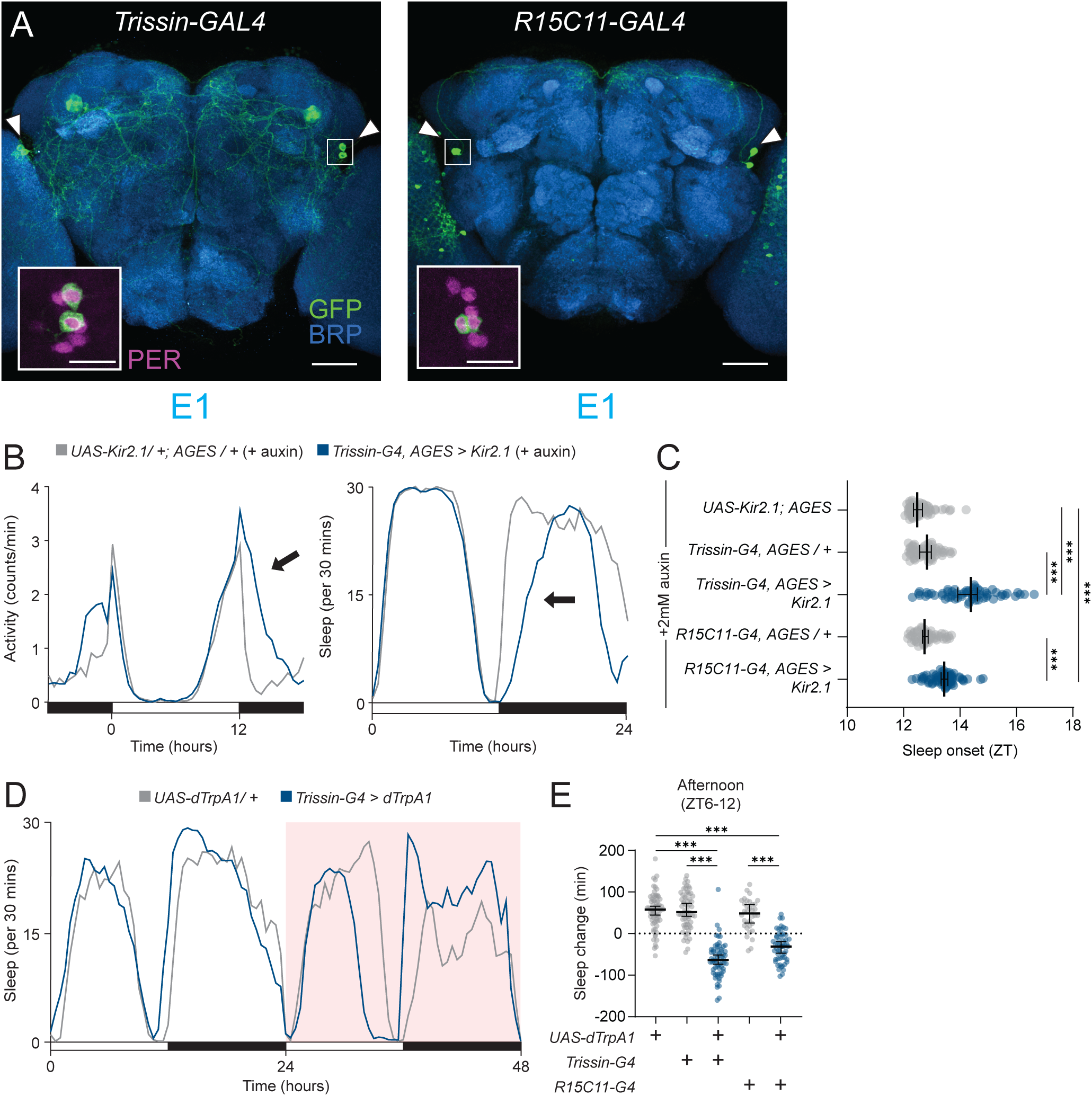
E1 neurons promote activity before dusk and sleep onset after dusk. **(A)** Immunostaining of E1 cell drivers. Whole-mount immunostaining of *Trissin-GAL4* (left) and *R15C11-GAL4* brains expressing *UAS-CD8::GFP* with anti-GFP (green), anti-BRP (blue), and anti-PER (magenta, inset) antibodies. E1 cells indicated by arrowheads. PER staining highlighted in solid white inset (LNds). Scale bars, 50 µm (main) and 15 µm (inset). **(B)** E1 neuron electrical silencing. Activity counts per min (left) and sleep amount per 30 min (right) profiles for 24 hrs for *Trissin-GAL4, AGES>UAS-Kir2.1* (blue) and *UAS-Kir2.1/+; AGES/+* (gray) flies in 12:12 LD plotted in 30 min bins. Arrows indicate delay in activity/sleep onset. Flies were kept on sucrose agar with 2 mM NAA during activity recording. **(C)** Time of sleep onset with E1 electrical silencing. Sleep onset time plotted by ZT for *Trissin-GAL4, AGES>Kir2.1* (blue, n=60), *R15C11-GAL4, AGES>Kir2.1* (blue, n=64), *Trissin-GAL4, AGES/+* (gray, n=59), *R15C11-GAL4, AGES/+* (gray, n=56), and *UAS-Kir2.1/+; AGES/+* (gray, n=56). Median ± 95% confidence interval is shown. Kruskal-Wallis with Dunn’s post hoc test. **(D)** E1 neuron 24 hr thermogenetic activation in LD. Sleep profile for 48 hrs for *Trissin-GAL4>UAS-dTrpA1* (blue) and *UAS-dTrpA1/+* (gray) flies in 12:12 LD plotted in 30 min bins. Red background indicates increased temperature (28°C), compared to 22°C baseline. **(E)** Afternoon sleep change with E1 neuron activation. Change in afternoon (ZT6-12) sleep amount with thermogenetic activation compared to baseline afternoon in *Trissin-GAL4>UAS-dTrpA1* (blue, n=63), *R15C11-GAL4>UAS-dTrpA1* (blue, n=55), *Trissin-GAL4/+* (gray, n=63), *R15C11-GAL4/+* (gray, n=34), and *UAS-dTrpA1/+* (gray, n=66) flies. Mean ± 95% confidence interval is shown. One-way ANOVA with Bonferroni post hoc test.

We next used AGES-induced expression of Kir2.1 to assess if conditionally silencing E1 neurons affects circadian locomotor activity. Strikingly, silencing E1 neurons in adult flies causes a significant delay in the rapid offset of activity after dusk at *zeitgeber* time (ZT) 12 (**Fig. 2B**). Flies exhibit long, consolidated bouts of sleep at night after dusk,^8–10^ but evening cells have not been previously shown to influence sleep timing. To address the possibility that E1 neurons affect the timing of sleep at night, we additionally measured the timing of the first sleep bout after dusk and found that E1 conditional silencing significantly delays sleep onset after dusk (**Fig. 2C**). Because fly clock neurons have previously been described as either “wake promoting” or “sleep promoting” (reviewed in Shafer and Keene, 2021),^34^ we suspected that E1 neurons might represent a novel “sleep promoting” cluster of clock neurons. To test whether E1 neurons induce sleep when activated, we used dTrpA1 to conditionally activate E1 neurons throughout the day. Surprisingly, activating E1 neurons produced time- and light-dependent effects on sleep. Instead of solely promoting sleep, E1 neuron activation induces almost no behavioral change in the morning, but a significant decrease in sleep in the afternoon (**Figs. 2D and 2E, S2E**). Perhaps most strikingly, the arousal-promoting effects of activating E1 neurons abruptly ceased at dusk, even though the E1 neurons were still being activated (**Fig. S2F**). Taken together, these findings suggest that E1 neurons play a multifaceted role in patterning behavior around dusk.

To further support that the observed effects on sleep and arousal were due to manipulating E1 neurons and not off-target neurons present in *Trissin-GAL4* or *R15C11-GAL4*, we tested a split-GAL4 combination that only labels E1 neurons in the central brain, *R18H11-AD; R78G02-DBD* (*E1 split-GAL4*, **Fig. S2G**). Using *E1 split-GAL4* to silence E1 neurons with Kir2.1 significantly delayed sleep onset (**Figs. S2H and S2I**). Moreover, thermogenetic activation of E1 neurons using *E1 split-GAL4* recapitulated the time- and light- dependent effects on sleep seen with *Trissin-GAL4* and *R15C11-GAL4* (**Figs. S2J and S2K**). These results using *E1 split-GAL4* confirm that the dramatic effects on behavior around dusk result from manipulating only the four E1 LNds.

### Dusk switches E1 behavioral output from arousal to sleep

Our 24-hr activation protocol suggested that the light-to-dark transition at dusk plays a key role in determining the behavioral output of the E1 neurons. However, to ensure the striking, light-dependent effect observed with this manipulation was not an artifact of 12-hr dTrpA1 activation before dusk, we activated E1 neurons for 3 hrs before and after dusk. Thermogenetic activation of E1 neurons using this regimen decreased sleep during the light phase and also advanced sleep onset time after lights-off (**Figs. 3A and 3B, S3A**). Notably, this latter phenotype is opposite to that seen with silencing E1 neurons, suggesting that E1 neural activity is responsible for timing sleep onset after dusk.

**Figure 3.**
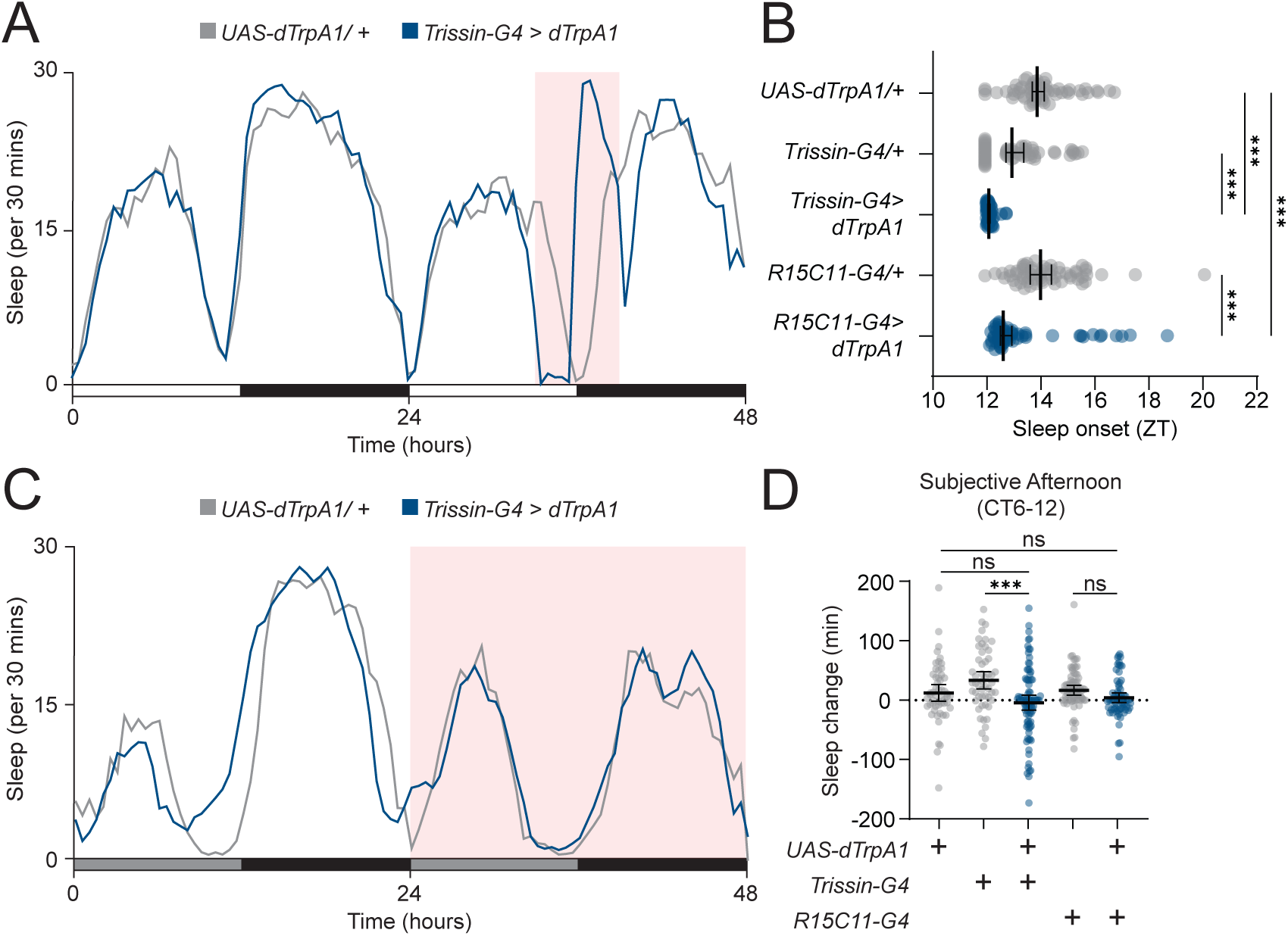
E1 neuron output is dependent on environmental light. **(A)** E1 neuron 6 hr dusk thermogenetic activation. Sleep profile for 48 hrs for *Trissin-GAL4>dTrpA1* (blue) and *UAS-dTrpA1/+* (gray) flies in 12:12 LD plotted in 30 min bins. Red background indicates increased temperature (29°C), compared to 22°C baseline. **(B)** Sleep onset with E1 neuron activation. Sleep onset time plotted by ZT for *Trissin-GAL4>UAS-dTrpA1* (blue, n=51), *R15C11-GAL4>UAS-dTrpA1* (blue, n=58), *Trissin-GAL4/+* (gray, n=56), *R15C11-GAL4/+* (gray, n=62), and *UAS-dTrpA1/+* (gray, n=59) flies. Median ± 95% confidence interval is shown. Kruskal-Wallis with Dunn’s post hoc test. **(C)** E1 neuron 24 hr thermogenetic activation in DD. Sleep profile for 48 hrs for *Trissin-GAL4>UAS-dTrpA1* (blue) and *UAS-dTrpA1/+* (gray) flies in DD plotted in 30 min bins. Red background indicates increased temperature (28°C), compared to 22°C baseline. Gray bars indicate subjective day in constant darkness. **(D)** Subjective afternoon sleep change with E1 neuron activation in DD. Change in afternoon (CT6-12) sleep amount compared to baseline afternoon for *Trissin-GAL4>UAS-dTrpA1* (blue, n=88), *R15C11-GAL4>UAS-dTrpA1* (blue, n=68), *Trissin-GAL4/+* (gray, n=52), *R15C11-GAL4/+* (gray, n=74), and *UAS-dTrpA1/+* (gray, n=54) flies. Mean ± 95% confidence interval is shown. One-way ANOVA with Bonferroni post hoc test.

After observing the sleep-advancing effects of E1 activation, we hypothesized that E1 neurons might simply promote arousal when activated in light conditions and sleep when activated in dark conditions. Alternatively, E1 neuron output might instead be sensitive to the transition from light to dark, meaning the sleep-promoting effects of E1 activation in the dark would not occur without a preceding light phase. To distinguish between these hypotheses, we activated E1 neurons for 24 hrs in constant darkness (DD), expecting to see an increase in sleep if our initial hypothesis was correct. Strikingly, E1 neuron activation in DD had no effect on locomotor behavior compared to controls (**Figs. 3C and 3D, S3B**), suggesting that E1 neuron output is sensitive to the transition from light to dark, rather than simply promoting arousal during the daytime and sleep during the nighttime. Overall, light appears to play two roles in determining the behavioral output of E1 neurons. First, our DD activation data suggest that light is a permissive signal that allows E1 neurons to influence behavior. Second, the data from our 6-hr activation around dusk suggest that the light-to-dark transition switches E1 behavioral output from promoting arousal to inducing sleep.

### Light input pathways act in parallel to gate E1-mediated behavior

Because light has a strong gating effect on E1 behavioral output in DD, we hypothesized that a specific light-sensing pathway might control that gating effect. Many clock neurons, including E1 neurons, can sense light in a cell-autonomous manner through the deep brain photoreceptor CRY.^35,36^ To test whether light sensation through CRY was gating E1 neuron behavioral output, we performed 24-hr dTrpA1 activation of E1 neurons in a *cry* null mutant background (*cry^02^*).^37^ If CRY were necessary for E1 output behavior, one would expect no change in behavior with E1 activation. However, similar to activation of E1 neurons in a wild-type background, activating E1 neurons in the *cry^02^* background in LD produced an afternoon reduction in sleep (**Figs. S3C and S3D**).

Outside of CRY, light can directly and indirectly interact with clock neurons via image-forming and non-image-forming visual pathways (reviewed in Yoshii *et al.*, 2016).^38^ Thus, we tested whether light sensation through the visual organs might be required for gating E1 behavioral output. To do this, we performed 24-hr activation of E1 neurons in LD in null mutants for the phospholipase C (PLC) required for visual transduction, *norpA* (*norpA^P24^*).^39^ Again, E1 neuron activation in the *norpA^P24^* background resulted in decreased sleep in the afternoon (**Figs. S3E and S3F**), suggesting that light sensation through visual pathways is also not, on its own, necessary for proper E1 behavioral output. Although the *norpA^P24^* allele should cause blindness in the compound eyes, ocelli, and H-B eyelets, some reports have suggested that photoreception in the Rhodopsin 6 (Rh6) expressing H-B eyelets is PLC-independent.^40,41^ Because the HB-eyelets form monosynaptic connections with clock network neurons,^18,42,43^ we further verified that the HB- eyelets were not necessary for gating E1 output behavior by thermogenetically activating E1 neurons for 24 hrs in an *rh6* null mutant background (*rh6*^1^).^44^ The *rh6* locus ends within 50 bases of the *trissin* locus, in which the *Trissin-2A-GAL4* is inserted, so we were only able to perform the *rh6*^1^ background experiments with *R15C11-GAL4*. As expected, *R15C11-GAL4>dTrpA1, rh6*^1^ flies also exhibited a significant reduction in sleep in the afternoon (**Figs. S3G and S3H**), confirming that the HB-eyelets are not necessary for gating E1 activation behavior. Because neither photoreception through CRY nor the visual system appear to be individually necessary for gating E1 output behavior, we suspect that these systems act in parallel to sense environmental light conditions and allow E1 activation to alter arousal and sleep behavior. Indeed, these systems appear to function in a redundant manner for other circadian behaviors, such as photoentrainment, photoperiod adjustment, and evening peak behavior in LD.^38,45^

### E1 neuron firing does not vary between dawn and dusk

Long-term Ca^2+^ imaging in clock network neurons has repeatedly shown that clusters of clock network neurons have daily rhythms of intracellular Ca^2+^.^11–13^ Additionally, electrophysiological analyses of multiple clusters of clock network neurons reveal daily changes in neural activity that correlate with the time of day when those cells influence behavior.^46–48^ Whether the electrical activity of LNd neurons varies across circadian time has not been previously addressed. However, these cells, as a group, do exhibit significantly higher levels of intracellular Ca^2+^ at or just before dusk.^11–13^ Because our silencing and activation experiments suggest that E1 LNds influence behavior around dusk, we hypothesized that their neural activity *in vivo* would be significantly higher at dusk compared to dawn. To test whether E1 neurons are more active at dusk (ZT12-13) or dawn (ZT0-1), we measured spiking frequency in E1 LNds *in vivo* using the voltage indicator Voltron2 (**Fig. 4A**).^49^ E1 neurons have symmetrical contra- and ipsilateral branching in the superior medial protocerebrum (SMP),^16,17^ allowing us to record changes in membrane potential and calculate spiking events from all four E1 LNds simultaneously in a single region of interest. Surprisingly, we did not detect a difference in LNd spiking rate between dawn and dusk (**Figs. 4B and 4C**), suggesting that changes in E1 activity are not necessary for their temporally restricted behavioral influence around dusk.

**Figure 4.**
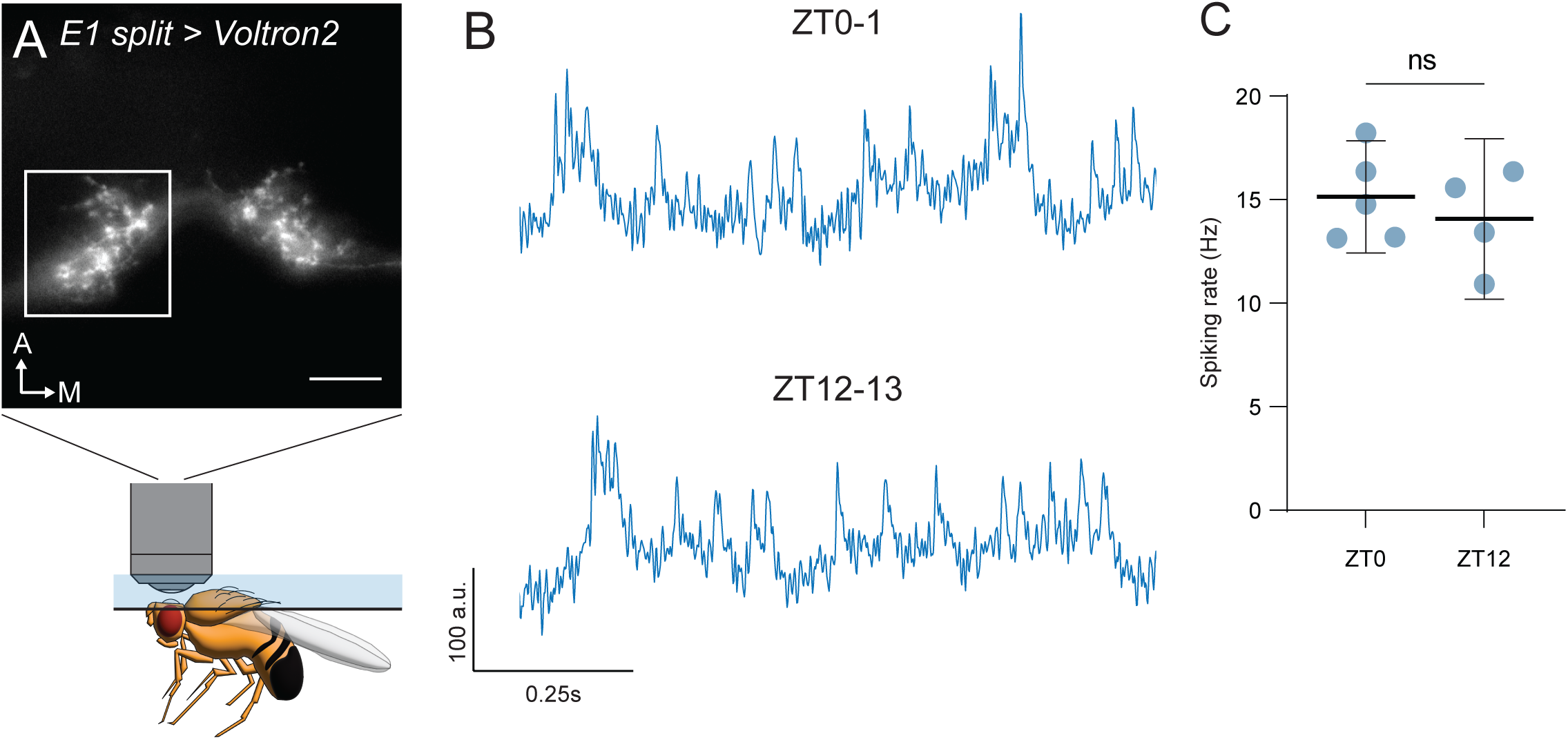
E1 neuron firing rate does not change between dawn and dusk. **(A)** Schematic of *in vivo* voltage imaging from E1 neurons. Still image of E1 processes in the superior medial protocerebrum (SMP) from a living *E1 split-GAL4>Voltron2* fly, illustrated below. White square represents approximate recording area. White arrows indicate anterior (top) and medial (right) directions. Scale bar, 25 µm. **(B)** Representative traces from dawn and dusk. Inverted fluorescence traces from *E1 split-GAL4>Voltron2* flies taken at dawn (top, ZT0-1) and dusk (bottom, ZT12-13). Scale bars represent change in fluorescence intensity (a.u., top) and time (seconds, bottom). **(C)** Spiking rate comparison between dawn and dusk recordings. Mean spiking rate for dawn (ZT0-1) and dusk (ZT12-13) recordings. Mean ± 95% confidence interval is shown. Student’s two-tailed t test.

### E1 neurons rely on a non-cell-autonomous clock to properly time behavioral output

Our E1 silencing and 24-hr activation experiments suggest that E1 behavioral output is temporally restricted to the evening, but our *in vivo* measurements of E1 activity suggest that their firing rate does not change between dawn and dusk. We next ensured that E1 behavioral output was being temporally gated by the circadian clock by performing thermogenetic activation of E1 neurons for 24 hrs in a *per^01^*background. Indeed, activating E1 neurons in a *per^01^* background causes an immediate loss of sleep starting at ZT0 that persists until ZT12 (**Figs. 5A and 5B**), suggesting that the circadian clock is responsible for temporally gating E1 activation-induced arousal.

**Figure 5.**
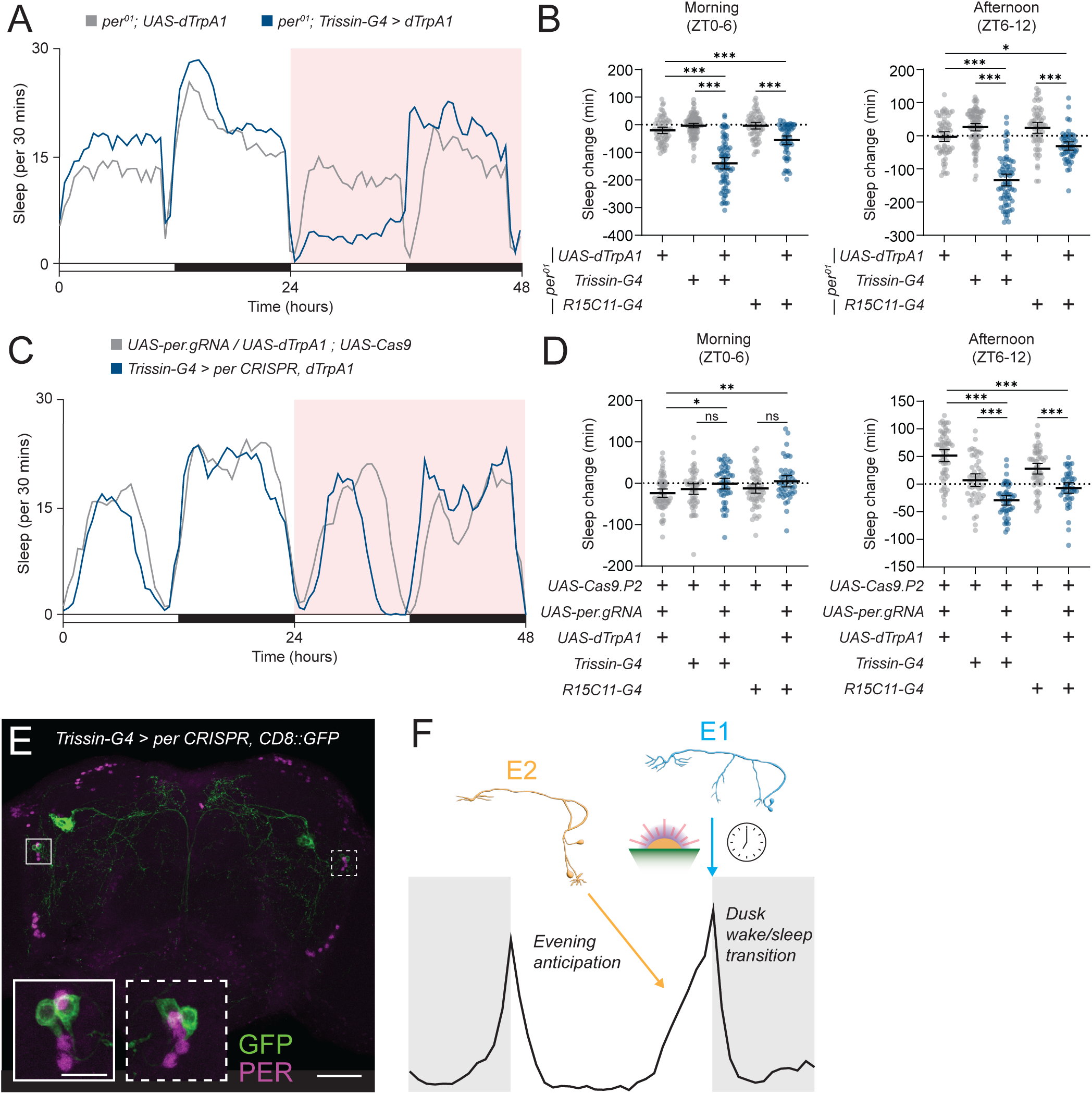
E1 behavioral output is dependent on the circadian clock in a non-cell autonomous manner. **(A)** E1 neuron 24 hr thermogenetic activation in *per* mutants. Sleep profile for 48 hrs for *per^01^;;Trissin-GAL4>UAS-dTrpA1* (blue) and *per^01^;UAS-dTrpA1/+* (gray) flies in 12:12 LD plotted in 30 min bins. Red background indicates increased temperature (28°C), compared to 22°C baseline. **(B)** Daytime sleep change with *per* mutant E1 neuron activation. Sleep change in the morning (left, ZT0-6) and afternoon (right, ZT6-12) compared to baseline day for *per^01^;;Trissin-GAL4>UAS-dTrpA1/+* (blue, n=71), *per^01^;;R15C11-GAL4>UAS-dTrpA1* (blue, n=55), *per^01^;;Trissin-GAL4/+* (gray, n=93), *per^01^;;R15C11-GAL4/+* (gray, n=60), and *per^01^;UAS-dTrpA1/+* (gray, n=62) flies. Mean ± 95% confidence interval is shown. One-way ANOVA with Bonferroni post hoc test. **(C)** E1 neuron 24 hr thermogenetic activation with tissue-specific *per* CRISPR knockout in LD. Sleep profile for 48 hrs for *Trissin-GAL4>UAS-dTrpA1, UAS-per.gRNA; UAS-Cas9.P2* (blue) and *UAS-dTrpA1, UAS-per.gRNA/+; UAS-Cas9.P2/+* (gray) flies in 12:12 LD plotted in 30 min bins. Red background indicates increased temperature (28°C), compared to 22°C baseline. **(D)** Daytime sleep change with E1 neuron activation and *per* knockout. Sleep change in the morning (left, ZT0-6) and afternoon (right, ZT6-12) compared to baseline day for *Trissin-GAL4>UAS-dTrpA1, UAS-per.gRNA; UAS-Cas9.P2* (blue, n=47), *R15C11-GAL4>dTrpA1, UAS-per.gRNA; UAS-Cas9.P2* (blue, n=45), *Trissin-GAL4>UAS-Cas9.P2* (gray, n=51), *R15C11-GAL4>UAS-Cas9.P2* (gray, n=57), and *UAS-dTrpA1, UAS-per.gRNA/+; UAS-Cas9.P2/+* (gray, n=63) flies. Mean ± 95% confidence interval is shown. One-way ANOVA with Bonferroni post hoc test. **(E)** Immunostaining of *per* tissue-specific knockout. Whole-mount immunostaining of a *Trissin-GAL4>UAS-CD8::GFP, UAS-per.gRNA; UAS-Cas9.P2* brain with anti GFP (green) and anti-PER (magenta). PER staining in GFP-positive LNds highlighted in insets. Scale bars, 50 µm (main) and 15 µm (inset). **(F)** Model describing the role of E1 and E2 neurons in shaping behavior around dusk. The cell- autonomous molecular clocks of E2 neurons (orange, left) mediate evening anticipation. At dusk, E1 neurons (cyan, right) facilitate the transition from wake to sleep in a light- and time- dependent manner.

Clock neurons are characterized by high-amplitude, rhythmic expression of core clock genes.^2^ Cell-autonomous clocks are thought to largely govern rhythmic output from these neurons with input from other clock neurons or environmental stimuli.^1,2,38,50,51^ Thus, we next asked whether *per* is specifically required in E1 neurons for timing their behavioral output. We performed tissue-specific CRISPR to knockout *per*^26^ in E1 neurons and then thermogenetically activated these cells for 24 hrs. Strikingly, this manipulation did not impair the timing of E1 output, showing no effect on behavior in the morning and a significant decrease in sleep in the afternoon (**Figs. 5C and 5D**). Using 3 *UAS* transgenes might dilute GAL4-UAS activity. Thus, to ensure that *per* tissue-specific CRISPR was effective in the presence of a 3^rd^ UAS-transgene, we performed PER immunostaining and found PER was largely eliminated in E1 neurons, even with the inclusion of an additional *UAS-mCD8::GFP* transgene (**Fig. 5E**). These data suggest that E1 neuron output is temporally restricted by a non-cell-autonomous clock that limits the behavioral influence of E1 neurons to the afternoon and evening.

It is possible that E1 behavioral output is timed by both the cell-autonomous clock and a non-cell-autonomous clock, the latter of which could compensate for the loss of the former. To test whether cell-autonomous clocks in E1 neurons were sufficient for timing E1 output behavior, we repeated the 24-hr E1 thermogenetic activation in the *per^01^* background while simultaneously rescuing *per* expression in E1 neurons using *UAS-per.16*.^52^ We found that restoring *per* expression in E1 neurons was not sufficient to properly pattern E1 output, as thermogenetic activation of E1 neurons in flies with *per* expression only in E1 clock neurons resulted in a significant reduction in sleep from ZT0-12, similar to that seen in *per^01^* null animals (**Figs. S4A and S4B**). These findings suggest that E1 neurons significantly, if not wholly, rely on a non-cell-autonomous clock to properly time their behavioral output.

## DISCUSSION

Since the first experiments almost two decades ago described the ability of evening cells to control evening anticipation in *Drosophila*,^3,4^ multiple lines of evidence have suggested that the evening lateral neurons are made up of distinct subclasses: E1, E2, and E3.^14–17,53,54^ However, to date there has not been a systematic attempt to define the behavioral roles of these subclasses in controlling circadian behavior. Here, our results show that between the CRY/PDFR-expressing E1 and E2 neurons, the cell-autonomous clocks of E2 neurons are primarily responsible for proper evening anticipation, the defining behavior of evening cells, and potently impact the phase of evening activity.

In this study, we also describe a behavioral role for the E1 LNds for the first time. E1 neuron activity appears to be important for flies to properly transition to sleep after dusk, and their behavioral output is strikingly reliant on the light-to-dark transition at dusk. However, we also found that their firing rate is not different between dawn and dusk. Additionally, their cell-autonomous clocks are dispensable for properly timing E1 behavioral output, suggesting they depend on timing information from other clock neurons to perform their functions at dusk (**Fig. 5F**).

### Cell-autonomous clocks, but not neural activity, in E2 neurons are required for evening anticipation

Although we showed that E2 neuron clocks are involved in evening anticipation, their neural activity appeared to be dispensable for this process, as neither silencing nor activation significantly affected evening anticipation. One reason E2 neural activity might be insulated from impacting anticipation is that E2 neurons receive direct synaptic input from the visual system and are strongly depolarized by light. Thus, if E2 neurons signal the evening phase of the day, then classical neurotransmission would be a poor method of communicating that information, as it would be confounded by firing induced by visual input. Instead, we hypothesize that the mechanism by which E2 neurons signal impending dusk is clock-dependent, but not activity-dependent. For example, a potential mechanism could be signaling via the constitutive secretory pathway, as the contents of these vesicles could be under circadian-dependent transcriptional/translational control, while their release could be Ca^2+^ independent.^55^

### E1 neurons control sleep timing in a light-dependent manner

To our knowledge, the E1 LNds represent the first example of a well-defined clock neuron subcluster that promotes both arousal and sleep, in this case in a light-dependent manner. Where does light act to regulate E1-mediated behavior? E1 neurons express CRY and are weakly depolarized by light through E2 neurons.^18^ However, we showed that CRY expression was not necessary for E1 activation-induced arousal. Moreover, thermogenetic activation of E1 neurons should more strongly depolarize these cells compared to light stimulation. Thus, we hypothesize that the gating effects of light on E1 neuron output occur downstream of E1 neurons.

Interestingly, we also showed that the visual system is not necessary for gating E1 output behavior. Because neither CRY nor visual input are required for E1-induced arousal, we suggest that parallel light pathways are involved in gating E1 output behavior. Multiple circuits could potentially receive external light information through parallel pathways. The ellipsoid body (EB), a neuropil closely associated with navigation and locomotion, contains CRY-expressing ring neurons and receives extensive synaptic input from the visual system.^11,56^ Additionally, tuberculobulbar (TuBu) neurons are part of the input pathway to the EB from the visual system and are also postsynaptic to CRY-positive clock neurons.^57–59^ It is possible that such parallel pathways exist to detect changes in a broad spectrum of light, as dusk changes both the brightness and composition of daylight.

### The clocks of E1 neurons are not required for their function at dusk

Clock neurons have traditionally been defined by their strong, rhythmic expression of core clock genes. It is generally assumed that both cell-autonomous expression of these genes and changes in spiking activity are necessary for a clock neuron subtype to function at a particular time of day.^2,12,13,50^ While E1 neurons impact behavior at dusk, we find that E1 LNds do not vary their firing rate between dawn and dusk. Although we initially predicted significant differences in E1 firing between these two timepoints, our behavioral experiments are consistent with these physiological results, suggesting that the cell-autonomous clocks of E1 LNds are of little importance for their behavioral output at dusk. Instead, we suspect that the rhythmicity of E1 neuron signaling is generated by a postsynaptic cluster of clock neurons (such as the dorsal neurons). For example, coupled networks of neuropeptidergic signaling are common in central clocks, including the mammalian SCN and *Drosophila* clock network.^14,60^ Thus, the dusk-specific actions of E1 neurons could be encoded by postsynaptic, clock-dependent synthesis of a neuropeptide receptor, such as TrissinR or sNPFR.^61,62^

### A neural circuit for sensing and reacting to dusk

One model for signaling time in clock neurons, both in the *Drosophila* clock network and the mammalian SCN, is that subsets of clock neurons exhibit daily rhythms of neural activity with different phase relationships throughout the day.^12,63,64^ While this mechanism is undoubtedly important for timekeeping in central clocks, even some neurons in the SCN do not exhibit rhythms in clock gene expression or electrical activity.^65–68^ We propose that E1 LNds illustrate an additional mechanism for timekeeping in the clock network in which some clock neurons signal the occurrence of an important *zeitgeber* landmark, like dusk. In other words, E1 neurons do not strictly function as “evening” neurons, but instead coordinate behaviors at dusk, a light-dependent time window.

We do not predict that E1 neurons sense and react to dusk on their own. Instead, we suggest that E1 neurons act as clock-accessory neurons that themselves are not rhythmic but still contribute to a larger circuit that together maintains daily, rhythmic behavior. In this case, E1 neurons prime the “dusk circuit” for transitioning from wakefulness to sleep, while other components of the dusk circuit signal the time and light conditions necessary to execute that behavior. Given the presence of functional parallels between the *Drosophila* clock network and the SCN, it is tempting to speculate that a similar circuit motif for sensing and reacting to twilight could also be present in mammals.

## ACKNOWLEDGEMENTS

We thank R. Allada and O. T. Shafer for kindly sharing *Drosophila* reagents. We also thank the Bloomington Stock Center for fly stocks (supported by NIH grant P40OD018537). We thank the Lavis lab for providing the JaneliaFluor dye for Voltron2 recording. We thank members of the Wu Lab for discussion. This work was supported by NIH grants F31NS117175 (M.P.B.), F32NS110183 (C.R.), R01NS079584 (M.N.W.), and R35NS122181 (M.N.W.).

## AUTHOR CONTRIBUTIONS

Conceptualization, MPB, MNW; Methodology, MPB, MFK, MNW; Software, MPB, MFK; Investigation, MPB, SV, IP, AGZ; Writing – Original Draft, MPB, MNW; Writing – Review & Editing, MPB, SV, IP, AGZ, CR, MFK, MNW; Resources, CR, MNW; Visualization, MPB; Supervision, MPB, MFK, MNW; Funding Acquisition, MPB, MNW.

## METHODS

### Transgenic fly strains

All GAL4, split-GAL4, and tissue-specific CRISPR lines were obtained from the Bloomington *Drosophila* stock center. Other flies were obtained as described in the key resources table.

### Immunostaining

Flies were collected and anaesthetized on ice between ZT22-2 for optimal PER visualization. Brains were removed in ice-cold phosphate buffered saline (PBS). After removing the surrounding fat bodies and trachea, the brains were fixed for 25 mins in a 4% solution of paraformaldehyde diluted in PBS at room temperature. Brains were then washed three times over 30 mins and blocked in a solution of 5% normal goat serum (NGS) 0.2% PBS-Triton X-100 (PBS-T) for 1-2 hrs at room temperature. Brains were then placed in a 5% NGS PBS-T solution containing primary antibody for two nights at 4°C. The following primary antibodies were used: chicken-anti-GFP (1:1000, ThermoFisher, A10262, RRID: AB_2534023), guinea pig-anti-PER^69^ (1:1000), and mouse-anti-BRP (1:20, DSHB nc82, RRID: AB_2314866). After primary antibody incubation, brains were washed in PBS-T four times over at least two hrs at room temperature. Brains were then placed in a 5% NGS PBS-T solution containing secondary antibody for one night at 4°C. Secondary antibodies were raised in goat against chicken, guinea pig, and mouse antisera and conjugated to Alexa Fluor 488, 568, or 647 (ThermoFisher A-11039, RRID: AB_2534096; ThermoFisher A-11075, RRID: AB_2534119; and ThermoFisher A-21235, RRID: AB_2535804) and were all diluted at a 1:1000 dilution. After secondary antibody incubation, brains were washed in PBS-T four times over at least 2 hrs in PBS-T at room temperature. Brains were then mounted on glass microscope slides using VectaShield Plus (Vector Labs) and size 1.5 coverslips.

### Confocal microscopy

Confocal images were acquired using an LSM710 microscope and Zen Black software (Zeiss). Pinhole aperture and slice thickness were adjusted to software recommendations for the given fluorophores and imaging parameters. Images shown are max intensity projections of multiple z slices. Central brain images to show driver specificity were taken using a 25x objective under oil immersion (“Plan-Neofluarℍ 25x/0.8). Zoomed in images of clock neuron GFP/PER expression were taken using a 63x objective under oil immersion.

### Behavioral measurements

All lines used in behavioral experiments were outcrossed at least four times to the *iso*^31^ background (BDSC 5905). Activity and sleep behavior were measured using the *Drosophila* Activity Monitoring system (Trikinetics) and the 5-minute of inactivity threshold for identifying sleep.^10^ Anticipation slope, sleep onset, and sleep amount were calculated using the Rethomics behavioral analysis package in R.^70^ Evening anticipation phase was calculated using PHASE.^71^ For all behavioral experiments, 2-4 day old male flies were loaded into sucrose locomotor tubes (5% sucrose, 2% agar), with the day of loading and following day discarded to allow for recovery from carbon dioxide anesthetization. For *UAS-dTrpA1* experiments, flies were raised at 23°C. Baseline behavioral data were collected at 22°C for these flies before being raised to 28°C for 24 hrs activation or 29°C for 6 hr activation. For DD *UAS-dTrpA1* experiments, one day of baseline behavior was recorded in LD (not shown) before releasing the animals into DD for one baseline day, then one day at 28°C. For all other experiments, flies were raised at 25°C and behavioral data were recorded at 25°C.

For conditional expression experiments using AGES,^30^ flies were raised on standard cornmeal molasses agar. Adult (2-3 day old) male flies were placed in NAA-containing (2 mM; PhytoTech Labs, N610) sucrose locomotor tubes for behavioral analysis.

### *In vivo* voltage imaging

Heterozygous male flies expressing *R18H11-AD; R78G02-DBD>UAS-Voltron2* (0-5 day old) were moved onto standard cornmeal molasses agar containing 0.5 mM all-trans retinal (Millipore-Sigma, R2500) 2-3 day before experiments. On the day of the recording, flies were taken out of their home incubator 2.5-3 hrs before the recording was to take place. Flies were fixed into a 3D-printed chamber with a chemically etched stainless steel shim bottom using UV-curable adhesive (Loctite AA 3972) on the head and thorax. The legs and proboscis were immobilized with beeswax. The imaging area was exposed by removing the cuticle, muscles, fat bodies, and trachea covering the dorsal part of the brain. The ocelli were kept intact and fixed behind the head against the cuticle. Dissection took place under oxygenated *Drosophila* physiological saline containing (in mM): NaCl, 103; KCl, 3; N-[Tris(hydroxymethyl)-methyl]-2-aminoethanesulfonic acid, 3; Trehalose dihydrate, 10; glucose, 10; sucrose, 2; NaHCO_3_, 26; NaH2PO_4_, 1; CaCl_2_, 1.5; MgCl_2_, 4. Following dissection, the chamber was filled with fresh dye-containing saline (1 µM JF552-HaloTag ligand), and the chamber then placed into a dark, humidified box for 1hr. The dye was washed out with three 5 min perfusions with saline at 1-1.5 mL/min. Each perfusion was followed by 10-15 mins of passive washing without perfusion. Widefield fluorescence imaging was performed using a fixed stage microscope (Olympus, BX51WI), a 60x water immersion objective (LUMPlanFl/IR, Olympus), and EMCCD camera (iXon Ultra, ANDOR). Excitation with a 530 nM LED (Luxeon) was controlled using a Mightex light source (BLS-1000-2) and BIOLED I/O controller (Mightex, BLS-019). Images were acquired at 849 Hz with 8x8 binning using Nikon imaging software (NIS-Elements).

Voltage imaging data were analyzed in MATLAB. Videos were motion-corrected using NoRMCorre,^72^ and ROIs were automatically selected using k-means clustering. After mean pixel intensity from the ROI was extracted for each frame, the trace was corrected for photobleaching by fitting a polynomial curve to the trace and subtracting the curve from the original trace. The trace was then filtered using a second order Butterworth filter with a 0.5 normalized cutoff frequency. Because the E1 neurons exhibited slow, oscillating subthreshold changes as well as spikes, the moving baseline fluorescence intensity was approximated using a 1-8 Hz bandpass filter. Spikes were automatically identified by finding local minima that were below two standard deviations from the estimated baseline fluorescence. These automatically detected spikes were manually inspected and corrected as needed. Spiking frequency was calculated by dividing the total number of identified spikes by the length of the recording.

### Quantification and statistical analysis

Statistical analyses were performed using Prism (GraphPad). For behavior experiments, pairwise comparisons were made between each experimental group (*GAL4>UAS*) and their respective genetic controls (*GAL4/+* or *UAS/+*). Differences between normally distributed data were calculated using one-way ANOVA with Bonferroni post-hoc test to correct for multiple comparisons. Differences in non-normally distributed data were calculated using Kruskal-Wallis with Dunn’s post-hoc test to correct for multiple comparisons. Sleep onset time, which was normally distributed in some experiments, but not in others due to the floor effect created by dusk, were all treated as though they were not normally distributed. For comparing two groups to each other, such as the spiking frequency in Voltron2 recordings, a two-tailed Student’s t test was used.

## Data availability

Code used for analyzing behavioral and voltage imaging data available at the following DOI: https://zenodo.org/badge/latestdoi/683218355.

---

**Figure S1.**
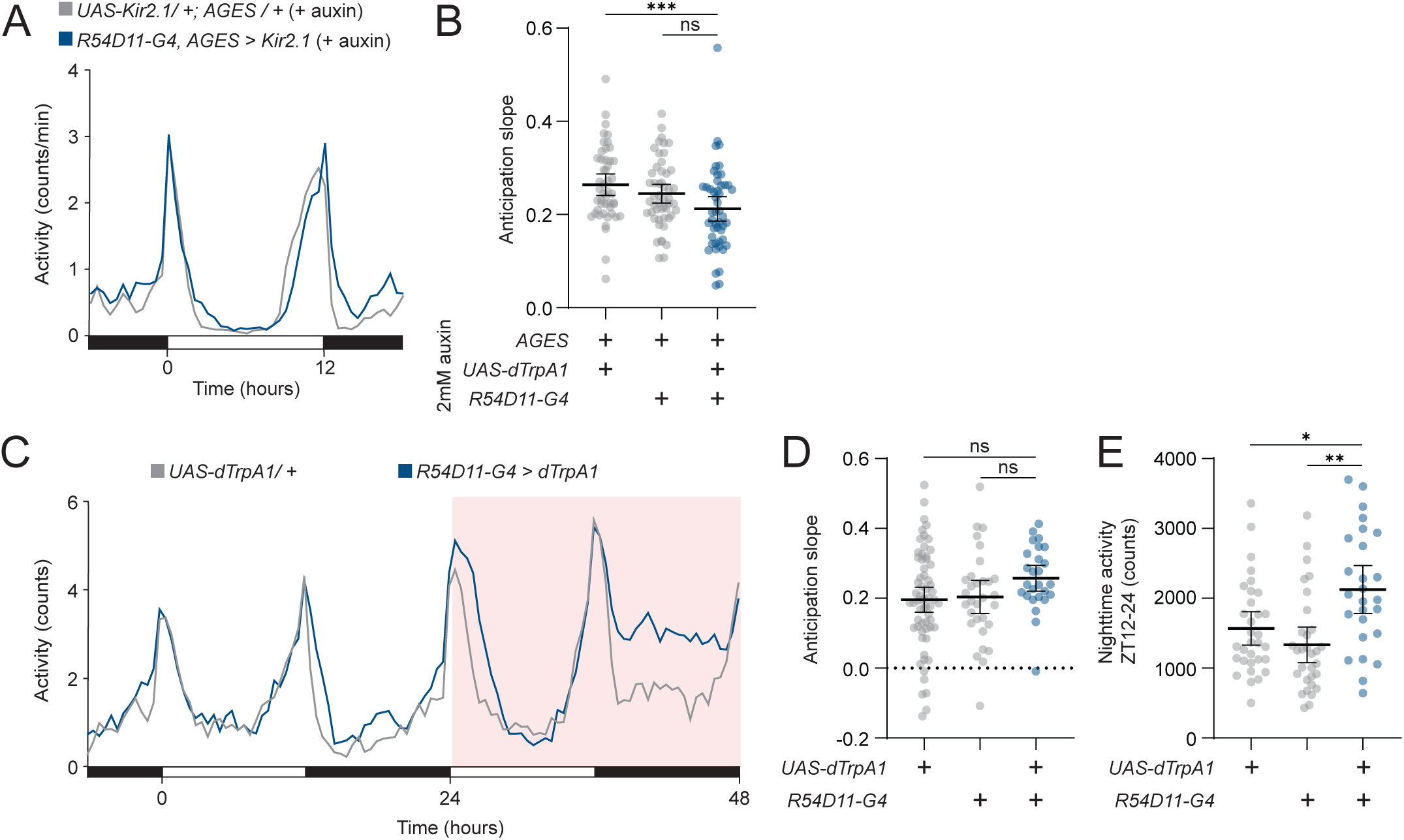
Altering the activity of E2 cells does not significantly affect evening anticipation, Related to Figure 1. **(A)** Conditional electrical silencing of E2 neurons. Activity counts per min profile for 24 hrs for *R54D11-GAL4, AGES>UAS-Kir2.1* (blue) and *UAS-Kir2.1/+; AGES/+* (gray) flies on 2 mM NAA sucrose agar in 12:12 LD plotted in 30 min bins. **(B)** Slope of evening anticipation for E2 conditional silencing. Average slope of evening anticipation (see Methods) plotted for *R54D11-GAL4, AGES>UAS-Kir2.1* (blue, n=49), *R54D11- GAL4, AGES/+* (gray, n=51), and *UAS-Kir2.1/+; AGES/+* (gray, n=48) flies. Mean ± 95% confidence interval is shown. One-way ANOVA with Bonferroni post hoc test. **(C)** E2 neuron conditional activation. Activity counts per min profile for 48 hrs for *R54D11-GAL4>UAS-dTrpA1* (blue) and *UAS-dTrpA1* (gray) flies in 12:12 LD plotted in 30 min bins. Red background indicates increased temperature (28°C), compared to 22°C baseline. **(D)** Slope of evening anticipation for E2 conditional activation. Average slope of evening anticipation (see Methods) plotted for activation day for *R54D11-GAL4>UAS-dTrpA1* (blue, n=26), *R54D11-GAL4/+* (gray, n=31), and *UAS-dTrpA1* (gray, n=32) flies. Mean ± 95% confidence interval is shown. One-way ANOVA with Bonferroni post hoc test. **(E)** Nighttime activity with E2 conditional activation. Average number of activity counts at night (ZT12-24) for *R54D11-GAL4>UAS-dTrpA1* flies (blue, n=26), *R54D11-GAL4/+* (gray, n=31), and *UAS-dTrpA1* (gray, n=32) flies. Mean ± 95% confidence interval is shown. One-way ANOVA with Bonferroni post hoc test.

**Figure S2.**
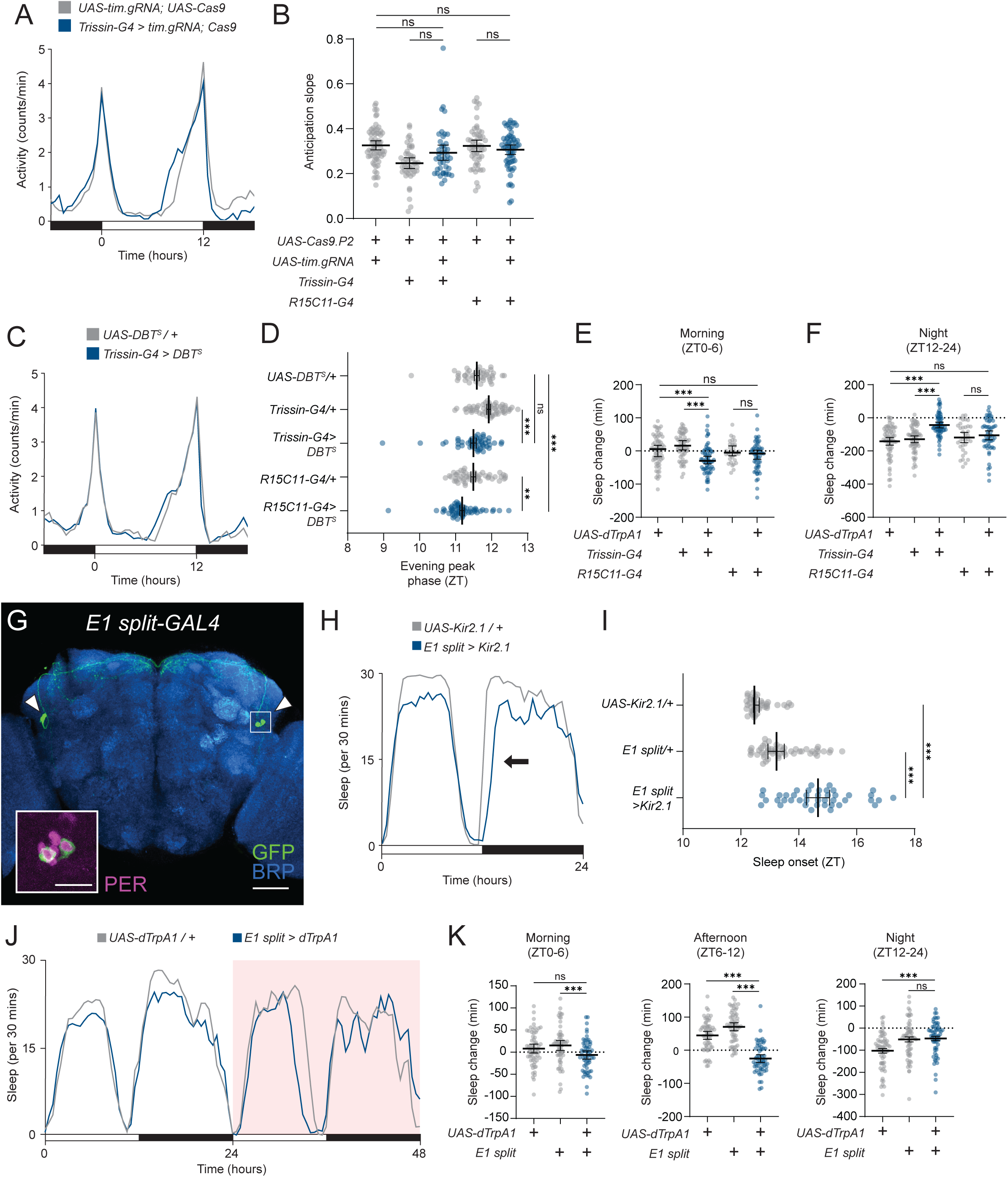
Electrical activity, not core clock function, in E1 neurons alters behavior around dusk, Related to Figure 2. **(A)** Tissue-specific knockout of *timeless* specifically in E1 neurons. Activity counts per min profile for 24 hrs for *Trissin-GAL4>UAS-tim.gRNA, UAS-Cas9.P2* (blue) and *UAS-tim.gRNA/+; UAS-Cas9.P2/+* (gray) flies in 12:12 LD plotted in 30 min bins. **(B)** Slope of evening anticipation for E1 neuron *tim* knockout. Average slope of evening anticipation (see Methods) for *Trissin-GAL4>UAS-tim.gRNA, UAS-Cas9.P2* (blue, n=43), *R15C11-GAL4>UAS-tim.gRNA, UAS-Cas9.P2* (blue, n=61), *Trissin-GAL4>UAS-Cas9.P2* (gray, n=46), *R15C11-GAL4>UAS-Cas9.P2* (gray, n=54), and *UAS-tim.gRNA/+; UAS-Cas9.P2/+* (gray, n=64) flies. One-way ANOVA with Bonferroni post hoc test. **(C)** DBT^S^ expression specifically in E1 neurons. Activity counts per min profile for 24 hrs for *Trissin-GAL4>UAS-DBT^S^* (blue) and *UAS-DBT^S^/+* (gray) flies in 12:12 LD plotted in 30 min bins. **(D)** Phase of evening anticipation for E1 neuron DBT^S^ expression. Average phase of evening anticipation for *Trissin-GAL4>UAS-DBT^S^* (blue, n=51), *R15C11-GAL4>UAS-DBT^S^* (blue, n=60), *Trissin-GAL4/+* (gray, n=57), *R15C11-GAL4/+* (gray, n=49), and *UAS-DBT^S^/+* (gray, n=41) flies. One-way ANOVA with Bonferroni post hoc test. **(E)** Morning sleep change with E1 neuron activation. Change in morning (ZT0-6) sleep amount with thermogenetic activation compared to baseline afternoon in *Trissin-GAL4>UAS-dTrpA1* (blue, n=63), *R15C11-GAL4>UAS-dTrpA1* (blue, n=55), *Trissin-GAL4/+* (gray, n=63), *R15C11-GAL4/+* (gray, n=34), and *UAS-dTrpA1/+* (gray, n=66). Same flies as Figs. 2D-E. Mean ± 95% confidence interval is shown. One-way ANOVA with Bonferroni post hoc test. **(F)** Nighttime sleep change with E1 neuron activation. Change in nighttime (ZT12-24) sleep amount with thermogenetic activation compared to baseline afternoon in *Trissin-GAL4>UAS-dTrpA1* (blue, n=63), *R15C11-GAL4>UAS-dTrpA1* (blue, n=55), *Trissin-GAL4/+* (gray, n=63), *R15C11-GAL4/+* (gray, n=34), and *UAS-dTrpA1/+* (gray, n=66) flies. Same flies as Figs. 2D-E. Mean ± 95% confidence interval is shown. One-way ANOVA with Bonferroni post hoc test. **(G)** Immunostaining of E1 split-GAL4 driver. Whole-mount immunostaining of an *R18H11-AD; R78G02-DBD* brain expressing *UAS-CD8::GFP* with anti-GFP (green), anti-BRP (blue), and anti-PER (magenta, inset) antibodies. E1 cells indicated by arrowheads. PER staining highlighted in solid white inset (LNds). Scale bars, 50 µm (main) and 15 µm (inset). **(H)** E1 split-GAL4 silencing behavioral trace. Sleep amount per 30 min (right) profiles for 24 hrs of *R18H11-AD; R78G02-DBD>UAS-Kir2.1* (blue) and *UAS-Kir2.1/+* (gray) flies in 12:12 LD plotted in 30 min bins. Arrow indicates delay in sleep onset. **(I)** Time of sleep onset with E1 split-GAL4 electrical silencing. Sleep onset time plotted by ZT for *R18H11-AD; R78G02-DBD>UAS-Kir2.1* (blue, n=42), *R18H11-AD; R78G02-DBD /+* (gray, n=51), and *UAS-Kir2.1/+* (gray, n=41) flies. Median ± 95% confidence interval is shown. Kruskal-Wallis with Dunn’s post hoc test. **(J)** E1 split-GAL4 24 hr thermogenetic activation in LD. Sleep profile for 48 hrs for *R18H11-AD; R78G02-DBD>UAS-dTrpA1* (blue) and *UAS-dTrpA1/+* (gray) flies in 12:12 LD plotted in 30 min bins. Red background indicates increased temperature (28°C), compared to 22°C baseline. **(K)** Daily sleep change with E1 split-GAL4 activation. Change in morning (ZT0-6, left), afternoon (ZT6-12, middle), and nighttime (ZT12-24, right) sleep amount with thermogenetic activation compared to baseline afternoon in *R18H11-AD; R78G02-DBD>UAS-dTrpA1* (blue, n=58), *R18H11-AD; R78G02-DBD/+* (gray, n=59), and *UAS-dTrpA1/+* (gray, n=61) flies. Mean ± 95% confidence interval is shown. One-way ANOVA with Bonferroni post hoc test.

**Figure S3.**
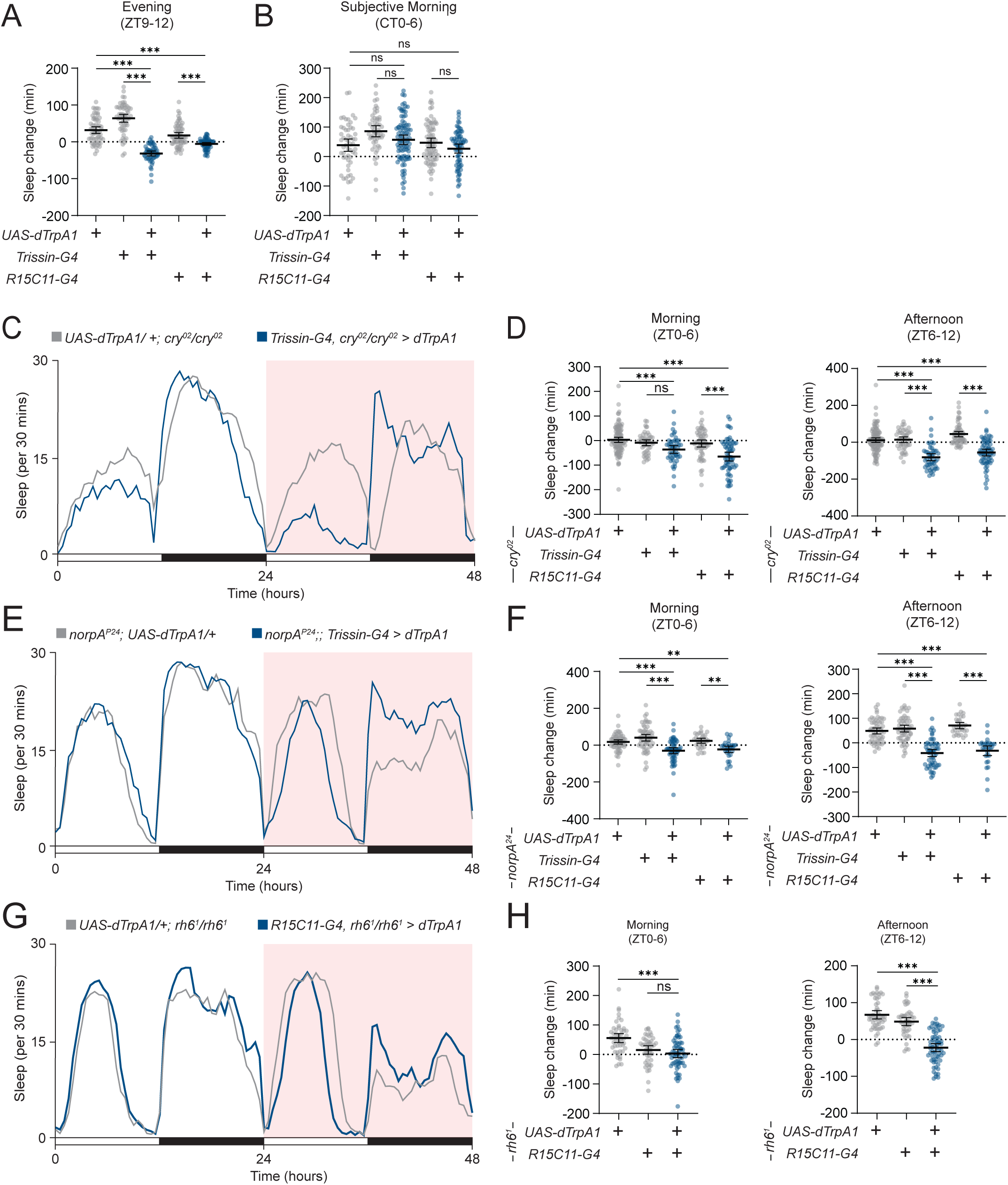
Light-dependent output of E1 neuron activation depends on parallel pathways, Related to Figure 3. **(A)** Evening sleep change with E1 neuron activation. Change in evening (ZT9-12) sleep amount with thermogenetic activation compared to baseline afternoon in *Trissin-GAL4>UAS-dTrpA1* (blue, n=51), *R15C11-GAL4>UAS-dTrpA1* (blue, n=58), *Trissin-GAL4/+* (gray, n=56), *R15C11-GAL4/+* (gray, n=63), and *UAS-dTrpA1/+* (gray, n=59) flies. Same flies as Figs. 3A-B. Mean ± 95% confidence interval is shown. One-way ANOVA with Bonferroni post hoc test. **(B)** Subjective morning sleep change with E1 neuron activation in DD. Change in morning (CT0-6) sleep amount with thermogenetic activation compared to baseline afternoon in *Trissin-GAL4>UAS-dTrpA1* (blue, n=88), *R15C11-GAL4>UAS-dTrpA1* (blue, n=68), *Trissin-GAL4/+* (gray, n=52), *R15C11-GAL4/+* (gray, n=74), and *UAS-dTrpA1/+* (gray, n=54) flies. Same flies as Figs. 3C-D. Mean ± 95% confidence interval is shown. One-way ANOVA with Bonferroni post hoc test. **(C)** E1 neuron 24 hr thermogenetic activation in LD in *cry* mutants. Sleep profile for 48 hrs for *Trissin-GAL4, cry^02^>UAS-dTrpA1; cry^02^*(blue) and *UAS-dTrpA1/+* (gray) flies in 12:12 LD plotted in 30 min bins. Red background indicates increased temperature (28°C), compared to 22°C baseline. **(D)** Daytime sleep change with E1 neuron activation in *cry* mutants. Change in morning (ZT0-6, left) and afternoon (ZT6-12, right) sleep amount with thermogenetic activation compared to baseline afternoon in *Trissin-GAL4, cry^02^>UAS-dTrpA1; cry^02^* (blue, n=47), *R15C11-GAL4, cry^02^>UAS-dTrpA1; cry^02^* (blue, n=63), *Trissin-GAL4, cry^02^/cry^02^* (gray, n=47), *R15C11-GAL4, cry^02^/cry^02^* (gray, n=60), and *UAS-dTrpA1/+*; *cry^02^/cry^02^*(gray, n=109) flies. Mean ± 95% confidence interval is shown. One-way ANOVA with Bonferroni post hoc test. **(E)** E1 neuron 24 hr thermogenetic activation in LD in *norpA* mutants. Sleep profile for 48 hrs for *norpA^P24^;; Trissin-GAL4>UAS-dTrpA1* (blue) and *norpA^P24^; UAS-dTrpA1/+* (gray) flies in 12:12 LD plotted in 30 min bins. Red background indicates increased temperature (28°C), compared to 22°C baseline. **(F)** Daytime sleep change with E1 neuron activation in *norpA* mutants. Change in morning (ZT0-6, left) and afternoon (ZT6-12, right) sleep amount with thermogenetic activation compared to baseline afternoon in *norpA^P24^;; Trissin-GAL4>UAS-dTrpA1* (blue, n=55), *norpA^P24^;; R15C11-GAL4>UAS-dTrpA1* (blue, n=30), *norpA^P24^;; Trissin-GAL4/+* (gray, n=58), *norpA^P24^;; R15C11-GAL4/+* (gray, n=31), and *norpA^P24^; UAS-dTrpA1/+* (gray, n=57) flies. Mean ± 95% confidence interval is shown. One-way ANOVA with Bonferroni post hoc test. **(G)** E1 neuron 24 hr thermogenetic activation in LD in *rh6* mutants. Sleep profile for 48 hrs for *R15C11-GAL4, rh6^1^>UAS-dTrpA1; rh6^1^* (blue) and *UAS-dTrpA1/+; rh6^1^/rh6^1^* (gray) flies in 12:12 LD plotted in 30 min bins. Red background indicates increased temperature (28°C), compared to 22°C baseline. **(H)** Daytime sleep change with E1 neuron activation in *rh6* mutants. Change in morning (ZT0-6, left) and afternoon (ZT6-12, right) sleep amount with thermogenetic activation compared to baseline afternoon in *R15C11-GAL4, rh6^1^>UAS-dTrpA1; rh6^1^* (blue, n=62), *R15C11-GAL4, rh6^1^/rh6^1^* (gray, n=45), and *UAS-dTrpA1/+; rh6^1^rh6^1^* (gray, n=49) flies. Mean ± 95% confidence interval is shown. One-way ANOVA with Bonferroni post hoc test.

**Figure S4.**
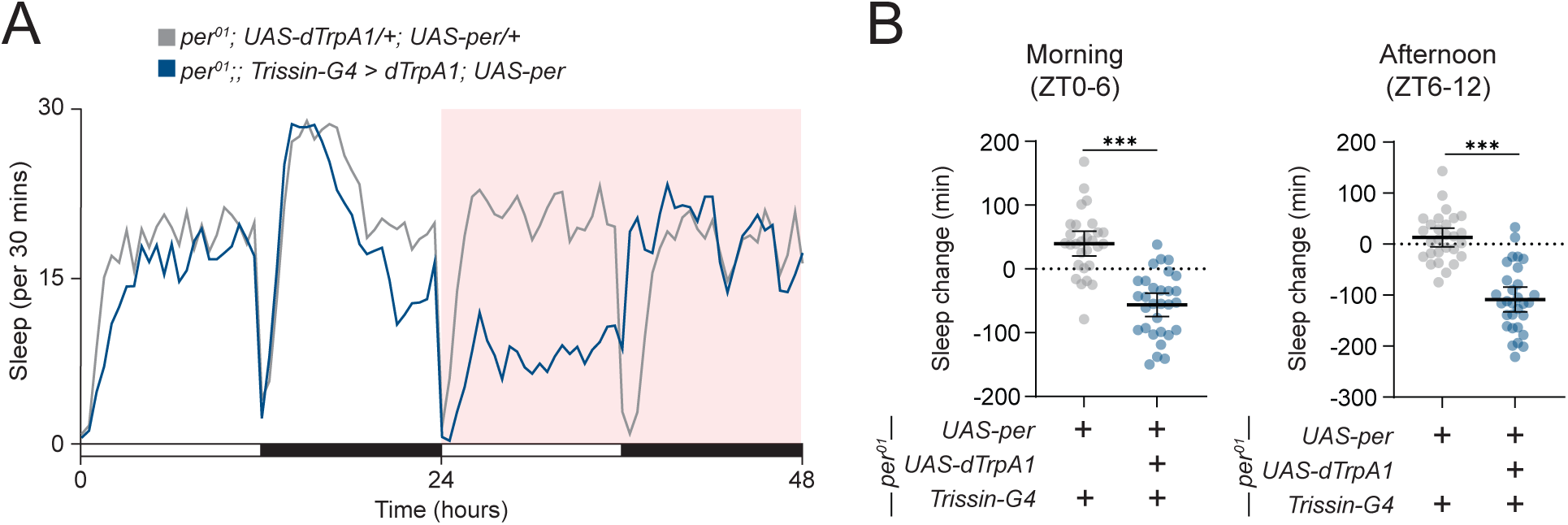
Rescue of *per* expression in E1 neurons is not sufficient to drive rhythmic output, Related to Figure 5. **(A)** E1 neuron 24 hr thermogenetic activation in *per* mutants with E1 *per* rescue. Sleep profile for 48 hrs of *per^01^;;Trissin-GAL4>UAS-dTrpA1; UAS-per^16^*(blue) and *per^01^;;Trissin-GAL4>UAS-per^16^* (gray) flies in 12:12 LD plotted in 30 min bins. Red background indicates increased temperature (28°C), compared to 22°C baseline. **(B)** Daytime sleep change with *per* mutant E1 neuron activation and E1 *per* rescue. Sleep change in the morning (left, ZT0-6) and afternoon (right, ZT6-12) compared to baseline day for *per^01^;;Trissin-GAL4>UAS-dTrpA1; UAS-per^16^* (blue, n=30) and *per^01^;;Trissin-GAL4>UAS-per^16^*(gray, n=28) flies. Mean ± 95% confidence interval is shown. One-way ANOVA with Bonferroni post hoc test.

## REFERENCES

1. Herzog, E.D. (2007). Neurons and networks in daily rhythms. Nature Reviews Neuroscience 8, 790–802. 10.1038/nrn2215.

2. Nitabach, M.N., and Taghert, P.H. (2008). Organization of the *Drosophila* Circadian Control Circuit. Current Biology 18, 84–93. 10.1016/j.cub.2007.11.061.

3. Grima, B., Chélot, E., Xia, R., and Rouyer, F. (2004). Morning and evening peaks of activity rely on different clock neurons of the *Drosophila* brain. Nature 431, 869–873. 10.1038/nature02935.

4. Stoleru, D., Peng, Y., Agosto, J., and Rosbash, M. (2004). Coupled oscillators control morning and evening locomotor behaviour of *Drosophila*. Nature 431, 862–868. 10.1038/nature02926.

5. Guo, F., Cerullo, I., Chen, X., and Rosbash, M. (2014). PDF neuron firing phase-shifts key circadian activity neurons in *Drosophila*. eLife 2014, 1–21. 10.7554/eLife.02780.

6. Yao, Z., Bennett, A.J., Clem, J.L., and Shafer, O.T. (2016). The *Drosophila* Clock Neuron Network Features Diverse Coupling Modes and Requires Network-wide Coherence for Robust Circadian Rhythms. Cell Reports 17, 2873–2881. 10.1016/j.celrep.2016.11.053.

7. Chatterjee, A., Lamaze, A., De, J., Mena, W., Chélot, E., Martin, B., Hardin, P., Kadener, S., Emery, P., and Rouyer, F. (2018). Reconfiguration of a Multi-oscillator Network by Light in the *Drosophila* Circadian Clock. Current Biology 28, 2007–2017.e4. 10.1016/j.cub.2018.04.064.

8. Rieger, D., Fraunholz, C., Popp, J., Bichler, D., Dittmann, R., and Helfrich-Förster, C. (2007). The Fruit Fly *Drosophila melanogaster* Favors Dim Light and Times Its Activity Peaks to Early Dawn and Late Dusk. J Biol Rhythms 22, 387–399. 10.1177/0748730407306198.

9. Hendricks, J.C., Finn, S.M., Panckeri, K.A., Chavkin, J., Williams, J.A., Sehgal, A., and Pack, A.I. (2000). Rest in *Drosophila* Is a Sleep-like State. Neuron 25, 129–138. 10.1016/S0896-6273(00)80877-6.

10. Shaw, P.J., Cirelli, C., Greenspan, R.J., and Tononi, G. (2002). Correlates of Sleep and Waking in *Drosophila melanogaster*. Science 287, 1834–1837. 10.1126/science.287.5459.1834.

11. Liang, X., Ho, M.C.W., Zhang, Y., Li, Y., Wu, M.N., Holy, T.E., and Taghert, P.H. (2019). Morning and Evening Circadian Pacemakers Independently Drive Premotor Centers via a Specific Dopamine Relay. Neuron, 843–857. 10.1016/j.neuron.2019.03.028.

12. Liang, X., Holy, T.E., and Taghert, P.H. (2016). Synchronous *Drosophila* circadian pacemakers display nonsynchronous Ca^2+^ rhythms in vivo. Science 351, 976–981. 10.1126/science.aad3997.

13. Liang, X., Holy, T.E., and Taghert, P.H. (2017). A Series of Suppressive Signals within the *Drosophila* Circadian Neural Circuit Generates Sequential Daily Outputs. Neuron 94, 1173–1189.e4. 10.1016/j.neuron.2017.05.007.

14. Yao, Z., and Shafer, O.T. (2014). The *Drosophila* Circadian Clock Is a Variably Coupled Network of Multiple Peptidergic Units. Science (New York, N.Y.) 343, 1516–1520. 10.1126/science.1251285.

15. Ma, D., Przybylski, D., Abruzzi, K.C., Schlichting, M., Li, Q., Long, X., and Rosbash, M. (2021). A transcriptomic taxonomy of *Drosophila* circadian neurons around the clock. eLife, 1–19. 10.1101/2020.09.15.297051.

16. Shafer, O.T., Gutierrez, G.J., Li, K., Mildenhall, A., Spira, D., Marty, J., Lazar, A.A., and M.P., F. (2022). Connectomic Analysis of the *Drosophila* Lateral Neuron Clock Cells Reveals the Synaptic Basis of Functional Pacemaker Classes. eLife 11, 1–29. 10.7554/eLife.79139.

17. Schubert, F.K., Hagedorn, N., Yoshii, T., Helfrich-Förster, C., and Rieger, D. (2018). Neuroanatomical details of the lateral neurons of *Drosophila melanogaster* support their functional role in the circadian system. Journal of Comparative Neurology 526, 1209–1231. 10.1002/cne.24406.

18. Li, M.T., Cao, L.H., Xiao, N., Tang, M., Deng, B., Yang, T., Yoshii, T., and Luo, D.G. (2018). Hub-organized parallel circuits of central circadian pacemaker neurons for visual photoentrainment in *Drosophila*. Nature Communications 9. 10.1038/s41467-018-06506-5.

19. Duhart, M., Herrero, A., Cruz, G.D., Ispizua, J.I., and Ceriani, M.F. (2020). Circadian Structural Plasticity Drives Remodeling of E Cell Output. Current Biology 30, 1–9. 10.1016/j.cub.2020.09.057.

20. Guo, F., Chen, X., and Rosbash, M. (2017). Temporal calcium profiling of specific circadian neurons in freely moving flies. Proceedings of the National Academy of Sciences of the United States of America 114, E8780–E8787. 10.1073/pnas.1706608114.

21. Bulthuis, N., Spontak, K.R., Kleeman, B., and Cavanaugh, D.J. (2019). Neuronal Activity in Non-LNv Clock Cells Is Required to Produce Free-Running Rest:Activity Rhythms in *Drosophila*. Journal of Biological Rhythms 34, 249–271. 10.1177/0748730419841468.

22. Siegmund, T., and Korge, G. (2001). Innervation of the ring gland of *Drosophila melanogaster*. J. Comp. Neurol. 431, 481–491. 10.1002/1096-9861(20010319)431:4<481::AID-CNE1084>3.0.CO;2-7.

23. Bahn, J.H., Lee, G., and Park, J.H. (2009). Comparative analysis of Pdf-mediated circadian behaviors between *Drosophila melanogaster* and *D. virilis*. Genetics 181, 965–975. 10.1534/genetics.108.099069.

24. Schlichting, M., Menegazzi, P., Lelito, K.R., Yao, Z., Buhl, E., Benetta, E.D., Bahle, A., Denike, J., Hodge, J.J., Helfrich-Förster, C., et al. (2016). A neural network underlying circadian entrainment and photoperiodic adjustment of sleep and activity in *Drosophila*. Journal of Neuroscience 36, 9084–9096. 10.1523/JNEUROSCI.0992-16.2016.

25. Yoshii, T., Hermann-Luibl, C., Kistenpfennig, C., Schmid, B., Tomioka, K., and Helfrich-Förster, C. (2015). Cryptochrome-dependent and -independent circadian entrainment circuits in *Drosophila*. Journal of Neuroscience 35, 6131–6141. 10.1523/JNEUROSCI.0070-15.2015.

26. Delventhal, R., O’Connor, R.M., Pantalia, M.M., Ulgherait, M., Kim, H.X., Basturk, M.K., Canman, J.C., and Shirasu-Hiza, M. (2019). Dissection of central clock function in *Drosophila* through cell-specific CRISPR-mediated clock gene disruption. eLife 8, 1–24. 10.7554/eLife.48308.

27. Stoleru, D., Peng, Y., Nawathean, P., and Rosbash, M. (2005). A resetting signal between *Drosophila* pacemakers synchronizes morning and evening activity. Nature 438, 238–242. 10.1038/nature04192.

28. Muskus, M.J., Preuss, F., Fan, J.-Y., Bjes, E.S., and Price, J.L. (2007). *Drosophila* DBT Lacking Protein Kinase Activity Produces Long-Period and Arrhythmic Circadian Behavioral and Molecular Rhythms. Molecular and Cellular Biology 27, 8049–8064. 10.1128/MCB.00680-07.

29. Baines, R.A., Uhler, J.P., Thompson, A., Sweeney, S.T., and Bate, M. (2001). Altered Electrical Properties in *Drosophila* Neurons Developing without Synaptic Transmission. The Journal of Neuroscience 21, 1523–1531. 10.1523/jneurosci.21-05-01523.2001.

30. McClure, C.D., Hassan, A., Aughey, G.N., Butt, K., Estacio-Gómez, A., Duggal, A., Sia, C.Y., Barber, A.F., and Southall, T.D. (2022). An auxin-inducible, GAL4-compatible, gene expression system for *Drosophila*. eLife 11. 10.7554/eLife.67598.

31. Hamada, F.N., Rosenzweig, M., Kang, K., Pulver, S.R., Ghezzi, A., Jegla, T.J., and Garrity, P.A. (2008). An internal thermal sensor controlling temperature preference in *Drosophila*. Nature 454, 217–220. 10.1038/nature07001.

32. Deng, B., Li, Q., Liu, X., Cao, Y., Li, B., Qian, Y., Xu, R., Mao, R., Zhou, E., Zhang, W., et al. (2019). Chemoconnectomics: Mapping Chemical Transmission in *Drosophila*. Neuron, 876–893. 10.1016/j.neuron.2019.01.045.

33. Sekiguchi, M., Inoue, K., Yang, T., Luo, D.G., and Yoshii, T. (2020). A Catalog of GAL4 Drivers for Labeling and Manipulating Circadian Clock Neurons in *Drosophila melanogaster*. Journal of Biological Rhythms 35, 207–213. 10.1177/0748730419895154.

34. Shafer, O.T., and Keene, A.C. (2021). The Regulation of *Drosophila* Sleep. Current Biology 31, R38–R49. 10.1016/j.cub.2020.10.082.

35. Ceriani, M.F., Darlington, T.K., Staknis, D., Más, P., Petti, A.A., Weitz, C.J., and Kay, S.A. (1999). Light-dependent sequestration of TIMELESS by CRYPTOCHROME. Science 285, 553–556. 10.1126/science.285.5427.553.

36. Fogle, K.J., Parson, K.G., Dahm, N. a, and Holmes, T.C. (2011). CRYPTOCHROME Is a Blue-Light Sensor That Regulates Neuronal Firing Rate. Science 331, 1409–1413. 10.1126/science.1199702.

37. Dolezelova, E., Dolezel, D., and Hall, J.C. (2007). Rhythm defects caused by newly engineered null mutations in *Drosophila*’s cryptochrome gene. Genetics 177, 329–345. 10.1534/genetics.107.076513.

38. Yoshii, T., Hermann-Luibl, C., and Helfrich-Föorster, C. (2016). Circadian light-input pathways in *Drosophila*. Communicative and Integrative Biology 9, 1–8. 10.1080/19420889.2015.1102805.

39. Pearn, M.T., Randall, L.L., Shortridge, R.D., Burg, M.G., and Pak, W.L. (1996). Molecular, Biochemical, and Electrophysiological Characterization of *Drosophila norpA* Mutants. Journal of Biological Chemistry 271, 4937–4945. 10.1074/jbc.271.9.4937.

40. Veleri, S., Brandes, C., Helfrich-Förster, C., Hall, J.C., and Stanewsky, R. (2003). A Self-Sustaining, Light-Entrainable Circadian Oscillator in the *Drosophila* Brain. Current Biology 13, 1758–1767. 10.1016/j.cub.2003.09.030.

41. Szular, J., Sehadova, H., Gentile, C., Szabo, G., Chou, W.-H., Britt, S.G., and Stanewsky, R. (2012). *Rhodopsin 5* – and *Rhodopsin 6* –Mediated Clock Synchronization in *Drosophila melanogaster* Is Independent of Retinal Phospholipase C-β Signaling. J Biol Rhythms 27, 25–36. 10.1177/0748730411431673.

42. Veleri, S., Rieger, D., Helfrich-Förster, C., and Stanewsky, R. (2007). Hofbauer-Buchner eyelet affects circadian photosensitivity and coordinates TIM and PER expression in *Drosophila* clock neurons. Journal of Biological Rhythms 22, 29–42. 10.1177/0748730406295754.

43. Helfrich-Förster, C., Edwards, T., Yasuyama, K., Wisotzki, B., Schneuwly, S., Stanewsky, R., Meinertzhagen, I.A., and Hofbauer, A. (2002). The extraretinal eyelet of *Drosophila*: Development, ultrastructure, and putative circadian function. Journal of Neuroscience 22, 9255–9266. 10.1523/jneurosci.22-21-09255.2002.

44. Cook, T., Pichaud, F., Sonneville, R., Papatsenko, D., and Desplan, C. (2003). Distinction between Color Photoreceptor Cell Fates Is Controlled by Prospero in *Drosophila*. Developmental Cell 4, 853–864. 10.1016/S1534-5807(03)00156-4.

45. Rieger, D., Stanewsky, R., and Helfrich-Förster, C. (2003). Cryptochrome, compound eyes, Hofbauer-Buchner eyelets, and ocelli play different roles in the entrainment and masking pathway of the locomotor activity rhythm in the fruit fly *Drosophila melanogaster*. Journal of Biological Rhythms 18, 377–391. 10.1177/0748730403256997.

46. Cao, G., and Nitabach, M.N. (2008). Circadian Control of Membrane Excitability in *Drosophila melanogaster* Lateral Ventral Clock Neurons. Journal of Neuroscience 28, 6493– 6501. 10.1523/JNEUROSCI.1503-08.2008.

47. Liu, S., Lamaze, A., Liu, Q., Tabuchi, M., Yang, Y., Fowler, M., Bharadwaj, R., Zhang, J., Bedont, J., Blackshaw, S., et al. (2014). WIDE AWAKE mediates the circadian timing of sleep onset. Neuron 82, 151–166. 10.1016/j.neuron.2014.01.040.

48. Tabuchi, M., Monaco, J.D., Duan, G., Bell, B., Liu, S., Liu, Q., Zhang, K., and Wu, M.N. (2018). Clock-Generated Temporal Codes Determine Synaptic Plasticity to Control Sleep. Cell 175, 1213–1227. 10.1016/j.cell.2018.09.016.

49. Abdelfattah, A.S., Zheng, J., Singh, A., Huang, Y.-C., Reep, D., Tsegaye, G., Tsang, A., Arthur, B.J., Rehorova, M., Olson, C.V.L., et al. (2023). Sensitivity optimization of a rhodopsin-based fluorescent voltage indicator. Neuron 111, 1547–1563.e9. 10.1016/j.neuron.2023.03.009.

50. King, A.N., and Sehgal, A. (2020). Molecular and circuit mechanisms mediating circadian clock output in the *Drosophila* brain. European Journal of Neuroscience 51, 268–281. 10.1111/ejn.14092.

51. Sehadova, H., Glaser, F.T., Gentile, C., Simoni, A., Giesecke, A., Albert, J.T., and Stanewsky, R. (2009). Temperature Entrainment of Drosophila’s Circadian Clock Involves the Gene nocte and Signaling from Peripheral Sensory Tissues to the Brain. Neuron 64, 251–266. 10.1016/j.neuron.2009.08.026.

52. Blanchardon, E., Grima, B., Klarsfeld, A., Chélot, E., Hardin, P.E., Préat, T., and Rouyer, F. (2001). Defining the role of *Drosophila* lateral neurons in the control of circadian rhythms in motor activity and eclosion by targeted genetic ablation and PERIOD protein overexpression. European Journal of Neuroscience 13, 871–888. 10.1046/j.0953-816X.2000.01450.x.

53. Im, S.H., and Taghert, P.H. (2010). PDF receptor expression reveals direct interactions between circadian oscillators in *Drosophila*. Journal of Comparative Neurology 518, 1925–1945. 10.1002/cne.22311.

54. Johard, H.A.D., Yoishii, T., Dircksen, H., Cusumano, P., Rouyer, F., Helfrich-Förster, C., and Nässel, D.R. (2009). Peptidergic clock neurons in *Drosophila*: Ion transport peptide and short neuropeptide F in subsets of dorsal and ventral lateral neurons. Journal of Comparative Neurology 516, 59–73. 10.1002/cne.22099.

55. Gondré-Lewis, M.C., Park, J.J., and Loh, Y.P. (2012). Cellular Mechanisms for the Biogenesis and Transport of Synaptic and Dense-Core Vesicles. In International Review of Cell and Molecular Biology (Elsevier), pp. 27–115. 10.1016/B978-0-12-394310-1.00002-3.

56. Scheffer, L., Xu, C.S., Januszewski, M., Lu, Z., Takemura, S., Hayworth, K., Huang, G., Shinomiya, K., Maitin-Shepard, J., Berg, S., et al. (2020). A connectome and analysis of the adult *Drosophila* central brain. eLife 9, 1–83. 10.1101/2020.04.07.030213.

57. Omoto, J.J., Keleş, M.F., Nguyen, B.C.M., Bolanos, C., Lovick, J.K., Frye, M.A., and Hartenstein, V. (2017). Visual Input to the *Drosophila* Central Complex by Developmentally and Functionally Distinct Neuronal Populations. Current Biology 27, 1098–1110. 10.1016/j.cub.2017.02.063.

58. Guo, F., Holla, M., Díaz, M., and Rosbash, M. (2018). A circadian output circuit controls sleep-wake arousal threshold in *Drosophila*. Neuron 100, 635. 10.1101/298067.

59. Lamaze, A., Kratschmer, P., Chen, K.-F., Lowe, S., and Jepson, J.E.C. (2018). A Wake-Promoting Circadian Output Circuit in *Drosophila*. Current Biology 28, 3098–3105. 10.1016/j.cub.2018.07.024.

60. Ono, D., Honma, K., and Honma, S. (2021). Roles of Neuropeptides, VIP and AVP, in the Mammalian Central Circadian Clock. Front. Neurosci. 15, 650154. 10.3389/fnins.2021.650154.

61. Ida, T., Takahashi, T., Tominaga, H., Sato, T., Kume, K., Yoshizawa-Kumagaye, K., Nishio, H., Kato, J., Murakami, N., Miyazato, M., et al. (2011). Identification of the endogenous cysteine-rich peptide trissin, a ligand for an orphan G protein-coupled receptor in *Drosophila*. Biochemical and Biophysical Research Communications 414, 44–48. 10.1016/j.bbrc.2011.09.018.

62. Feng, G., Reale, V., Chatwin, H., Kennedy, K., Venard, R., Ericsson, C., Yu, K., Evans, P.D., and Hall, L.M. (2003). Functional characterization of a neuropeptide F-like receptor from *Drosophila melanogaster*. Eur J Neurosci 18, 227–238. 10.1046/j.1460-9568.2003.02719.x.

63. Evans, J.A. (2016). Collective timekeeping among cells of the master circadian clock. Journal of Endocrinology 230, R27–R49. 10.1530/JOE-16-0054.

64. Welsh, D.K., Logothetis, D.E., Meister, M., and Reppert, S.M. (1995). Individual neurons dissociated from rat suprachiasmatic nucleus express independently phased circadian firing rhythms. Neuron 14, 697–706. 10.1016/0896-6273(95)90214-7.

65. Jiao, Y.-Y., Lee, T.M., and Rusak, B. (1999). Photic responses of suprachiasmatic area neurons in diurnal degus (*Octodon degus*) and nocturnal rats (*Rattus norvegicus*). Brain Research 817, 93–103. 10.1016/s0006-8993(98)01218-9.

66. Hamada, T., LeSauter, J., Venuti, J.M., and Silver, R. (2001). Expression of *Period* Genes: Rhythmic and Nonrhythmic Compartments of the Suprachiasmatic Nucleus Pacemaker. J. Neurosci. 21, 7742–7750. 10.1523/JNEUROSCI.21-19-07742.2001.

67. Jobst, E.E., and Allen, C.N. (2002). Calbindin neurons in the hamster suprachiasmatic nucleus do not exhibit a circadian variation in spontaneous firing rate: Firing rate of calbindin neurons in hamster SCN. European Journal of Neuroscience 16, 2469–2474. 10.1046/j.1460-9568.2002.02309.x.

68. Karatsoreos, I.N., Yan, L., LeSauter, J., and Silver, R. (2004). Phenotype Matters: Identification of Light-Responsive Cells in the Mouse Suprachiasmatic Nucleus. J. Neurosci. 24, 68–75. 10.1523/JNEUROSCI.1666-03.2004.

69. Sidote, D., Majercak, J., Parikh, V., and Edery, I. (1998). Differential Effects of Light and Heat on the *Drosophila* Circadian Clock Proteins PER and TIM. Molecular and Cellular Biology 18, 2004–2013. 10.1128/MCB.18.4.2004.

70. Geissmann, Q., Rodriguez, L.G., Beckwith, E.J., and Gilestro, G.F. (2019). Rethomics: An R framework to analyse high-throughput behavioural data. PLoS ONE 14, 1–19. 10.1371/journal.pone.0209331.

71. Persons, J.L., Abhilash, L., Lopatkin, A.J., Roelofs, A., Bell, E.V., Fernandez, M.P., and Shafer, O.T. (2022). PHASE: An Open-Source Program for the Analysis of *Drosophila* Phase, Activity, and Sleep Under Entrainment. J Biol Rhythms 37, 455–467. 10.1177/07487304221093114.

72. Pnevmatikakis, E.A., and Giovannucci, A. (2017). NoRMCorre: An online algorithm for piecewise rigid motion correction of calcium imaging data. Journal of Neuroscience Methods 291, 83–94. 10.1016/j.jneumeth.2017.07.031.

